# LRP8 is a Receptor for Yellow Fever Virus

**DOI:** 10.1101/2025.10.28.685229

**Authors:** Miao Mei, Yang Yang, Zihan Zhang, Yue Yin, Jiali Tan, Chao Jiang, Yu Gao, Zhaoyang Wang, Donghong Wang, Yajing Li, Yingyi Cong, Zhiyuan Zhang, Yousong Peng, Wenjie Tan, Jiandong Li, Li Li, Hanxue Wang, Ren Lang, Qiang He, Zihou Deng, Xiaojie Huang, Cao Shan, Yonghong Zhang, Gong Cheng, Xu Tan

## Abstract

Yellow fever virus (YFV), an arbovirus causing substantial human morbidity and mortality, was the first human virus discovered over a century ago. The live-attenuated 17D vaccine is among the most successful vaccines in medicine. Despite the importance of YFV, its receptor has remained unknown. Here, we performed a barcoded, genome-wide human ORF library screen and identified LRP8 (also named APOER2) as a receptor for YFV. We show that LRP8 expression specifically boosts YFV infection in cell lines by promoting entry. AAV-mediated expression of human LRP8 in mouse liver aggravates infection and pathology. LRP8 knockdown abolishes YFV infection in brain cells, primary human hepatocytes, and notably in mosquitoes. Biochemically, LRP8 directly interacts with YFV particles via the viral envelope protein. This function of LRP8 is conserved across species, particularly in mosquitoes and primates. A soluble LRP8 decoy protein can block YFV infection in vitro and in mice, providing a potential therapeutic or prophylactic strategy. Our findings provide groundwork for understanding YFV entry, tropism, and pathogenesis, and may enable development of novel therapeutics to treat YFV infection.

## Introduction

Yellow fever virus (YFV), the first human virus ever discovered, is among the deadliest human pathogens and has shaped human history^1–3^. Owing to the landmark discovery in 1900^4^ that mosquitoes transmit YFV and subsequent public-health efforts—particularly development and wide application of the 17D vaccine—Yellow fever (YF) is no longer considered epidemic in most countries today^5^. However, the sylvatic transmission cycle of YFV, namely transmission between non-human primates via forest mosquitoes, makes elimination highly improbable. YF remains a significant public-health concern in tropical regions of South America and Africa, with an estimated 84,000–170,000 severe cases and 29,000–60,000 deaths per year^6^. Recent outbreaks occurred in Brazil, the Democratic Republic of the Congo, Angola, Uganda, and Nigeria, with spillover cases in Asia ^7^. YFV infection causes hemorrhagic fever characterized by hepatitis and jaundice. YFV infects the liver and can induce disseminated intravascular coagulation, producing hemorrhagic manifestations^8^. In late stages, the central nervous system may be affected together with multiple organ failure preceding death^8^.

The attenuated 17D vaccine, developed in the 1930s, has been administered to over 500 million people, greatly reducing the burden of YFV for decades. However, vaccine production constraints during emergent outbreaks hinder rapid, comprehensive immunization^7,9^. In addition, rare serious adverse events such as Yellow Fever Vaccine-Associated Neurotropic Disease (YEL-AND) and Yellow Fever Vaccine-Associated Viscerotropic Disease (YEL-AVD) suggest directions for improvement in next-generation vaccines^10,11^. The 17D vaccine was obtained by serial passage of the Asibi strain in mouse brain and chicken embryos (>176 passages), resulting in reduced neuro- and viscerotropism. Sequencing pinpointed 20 amino-acid differences and four additional nucleotide differences between 17D and Asibi that underlie reduced virulence^12^. Among the 20 amino-acid changes, eight reside in the envelope (E) protein, which determines tropism by engaging cellular receptors and mediating entry.^12–16^.

A better understanding of the YFV life cycle would clarify tropism, pathogenesis, and the mechanism of 17D attenuation. Currently, the identity of the specific cellular receptor for YF remains unknown, hampering a molecular understanding of entry and virulence^17^. Here, we describe a genome-wide gene-overexpression screen that identifies LRP8 (also named APOER2) as a receptor for YFV. We show that LRP8 is expressed in both liver and brain, illuminating the tissue tropism of YF. Our results also highlight the effectiveness of overexpression screening in identifying viral receptors that have resisted gene-downregulation approaches such as CRISPR-Cas9 and RNAi.

### Genome-wide overexpression screen to identify LRP8 as a potential YFV receptor

To uncover the long-sought YFV receptor, we applied a barcoded ORFeome library covering more than 15,000 human genes in a lentivector system driven by a CMV promoter to facilitate stable expression in mammalian cells^18,19^. We performed the screen in HEK293T cells, which have intrinsically low susceptibility to YFV. Cells were transduced with the ORFeome library at low MOI (0.1) to ensure single-ORF expression per cell. After puromycin selection, cells were infected with a recombinant YFV reporter (GFP) virus (**Fig.1a**)^20^. GFP expression enabled flow-cytometric sorting to enrich infected cells. GFP-positive and GFP-negative populations were separately processed for genomic DNA extraction, followed by PCR amplification of the barcode region adjacent to the ORF C terminus. Deep sequencing quantified the relative abundance of each ORF in the two populations. The screen was performed twice, each in triplicate. Low-density lipoprotein receptor-related protein 8 (LRP8) consistently ranked among the top candidates enriched in GFP-positive cells (**Fig.1b and Supplementary Data Set**). LRP8 (ApoER2) belongs to the low-density lipoprotein receptor (LDLR) family ^21^. It is known to be expressed in the brain and plays crucial roles in neuronal development and synaptic plasticity ^22,23^. Previously, LRP8 together with VLDLR, another LDLR family member, was identified as an entry receptor for certain alphaviruses including Semliki Forest virus, eastern equine encephalitis virus, and Sindbis virus^24^. However, to the best of our knowledge, no study implicated LRP8 in flavivirus infection.

Overexpression of LRP8 dramatically enhanced infection by both 17D pseudovirus, but not Zika pseudovirus, as shown by image quantification (**Fig. 1c and 1d**). Plaque assays showed that LRP8 significantly boosted infection of YFV-17D live virus (**Fig. 1e**). Moreover, three major LRP8 isoforms showed similar activity in promoting YF infection (**Extended Fig. 1a-1c**). In contrast, overexpression of TIM1, a phosphoserine-binding protein, facilitated Zika virus infection^25^ and had much weaker effect on YFV infection (**Fig. 1c and 1d, Extended Fig. 1e and 1f**). LDLR, the prototypic member of the LDLR family, showed little effect on YFV infection in our ORFeome screen and validation experiments (**Fig. 1b, 1c and 1d**). Because the ligand-binding domain (LBD) of LRP8 mediates engagement of endogenous ligand Reelin and alphavirus envelopes^12,26^, we generated two LRP8 constructs: an LBD deletion (ΔLBD) and an EGF-domain deletion (ΔEGF) control. Ectopic expression of LRP8-ΔLBD lost the ability to promote YFV infection, whereas LRP8-ΔEGF retained activity (**Extended Fig. 1a, 1b and 1d**), supporting a requirement for the LBD. It is worth mentioning that LRP8-ΔLBD also has much lower protein expression than those of LRP8-ΔEGF and full length protein, suggesting multiple roles of LBD. We also expressed LRP8 in K562 cells, a line with very low levels of endogenous LRP8^24^ and showed that it also promoted YFV-17D infection (**Extended Fig. 2**). Beyond the 17D vaccine strain, we tested BJ01, a clinical isolate of the Angola genotype derived from a rare spillover case in China^27^. YFV-BJ01 showed enhanced infection as 17D in LRP8-overexpressing cells (**Fig. 1e, 1f and 1g, Extended Fig 3a, 3b**). BJ01 pseudovirus infection was likewise enhanced by LRP8 (**Extended Fig. 3c**). Pseudotyped particles bearing E proteins from Dengue virus, Japanese encephalitis virus, West Nile virus, Sepik, or Wesselsbron viruses, the latter two the closest to YFV based on genome analysis^28^, did not exhibit increased infection in LRP8-overexpressing cells (**Fig. 1h**). Overall, LRP8 promotes YFV entry in a rather specific fashion among the viruses tested.

**Figure 1.**
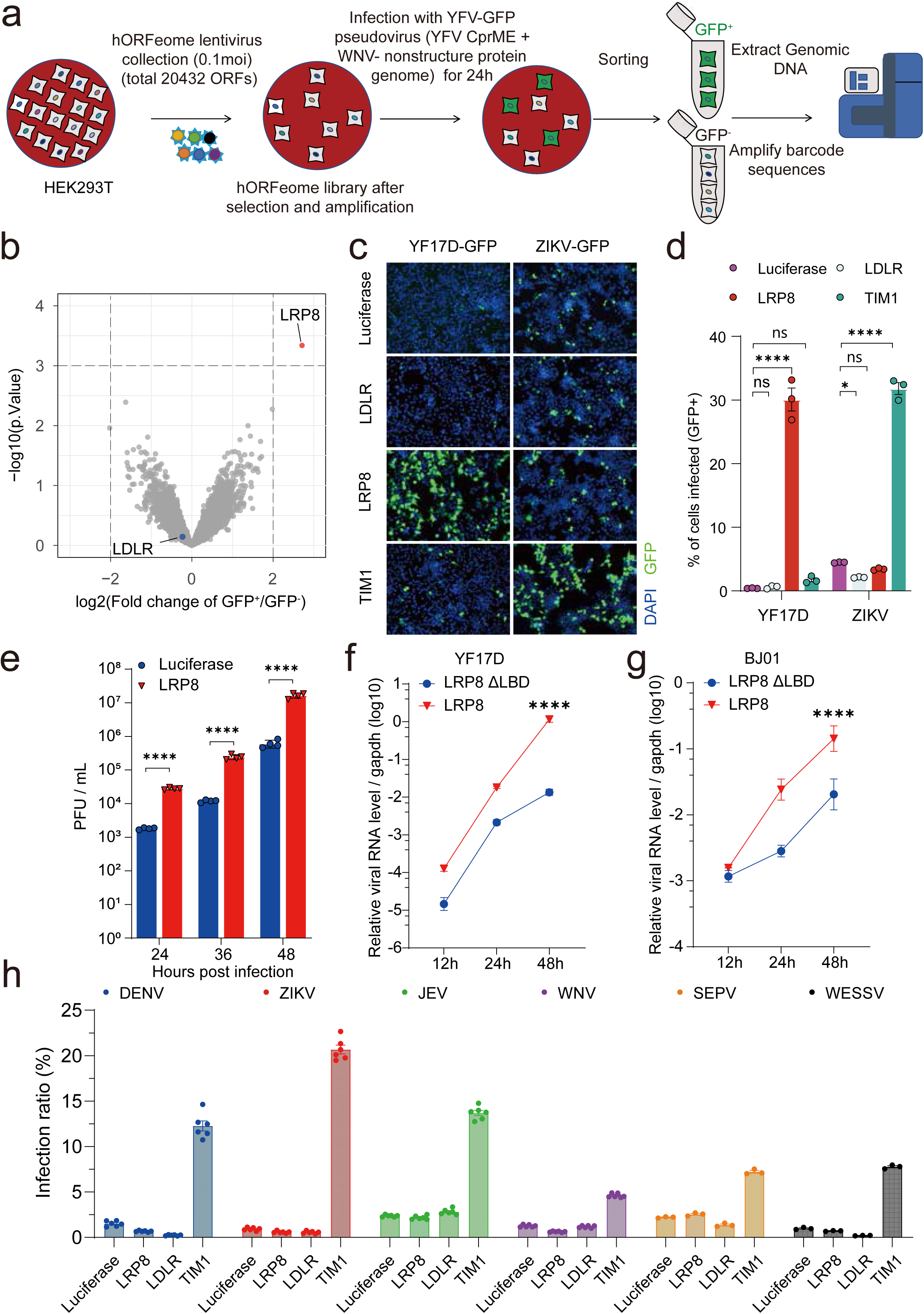
Human genome-wide ORFeome library screening identifies LRP8 as a YFV receptor. **a,** Schematic of the workflow of the screen. **b,** Deep-sequencing analysis of the screen performed with YF17D-GFP in HEK293T cells stably expressing the human ORFeome library. The x-axis shows log2 fold change of a gene’s read counts in GFP+ versus GFP− cells; the y-axis shows −log10 p-values. **c-e,** Luciferase, LDLR, LRP8, or TIM1 were expressed in HEK293T cells and infected with YF17D, ZIKV, Dengue virus 2, WNV, JEV, SEPV or WESSV pseudoviruses. Blue and green indicate DAPI and GFP, respectively. **d**, Quantification of **c**. **e**, Supernatants from HEK293T overexpressing luciferase or LRP8 infected by live YF17D virus at 48 hpi were collected and the viral titers were determined using palque assay in BHK21 cells. **f-g**, Ectopic LRP8 expression enhanced infection of YF17D (MOI 0.1) (**e**) and BJ01 (MOI 0.01) (**f**) shown in a time course. Viral RNA was normalized to GAPDH. h. HEK293T cells were transfected with the indicated genes were infected with the indicated pseudoviruses for 48h before being analyzed by FACS. Statistical analysis was performed using one-way analysis of variance (ANOVA) with Dunnett’s multiple comparisons test(d), two-tailed unpaired t-tests (e, f, g).The statistics shown are Mean ± SEM. *: P < 0.05; **: P < 0.01; ***: P < 0.001; ****: P < 0.0001, ns, not significant. The replicate in the figure indicates biological replicates, the experiments were repeated at least three times.

We also generated U87-MG *LRP8* knockout (KO) cells, a glioma-origin cell line. The KO was validated by FACS analysis and Western blots (**Extended Fig. 4a and Fig. 2a**) The *LRP8*-KO cells showed profound decreases in infection by YFV 17D (**Fig. 2a, 2b and 2c**) and the clinical strains BJ01 and Asibi (**Fig. 2d**, **2e and Extended Fig. 4b**). Importantly, this decrease was reversed by ectopic expression of sgRNA-resistant LRP8 constructs with synonymous mutations in sgRNA sites, ruling out CRISPR off-target effects (**Fig. 2f, 2g and 2h, Extended Fig. 4c, 4d and 4e**). Consistently, knockout of LRP8 had no effect on Zika virus infection (**Fig. 2a, 2b and 2c**). Moreover, an antibody targeting the extracellular domain of LRP8 (sc-293472) blocked YFV but not Zika virus, further supporting a cell-surface receptor role (**Fig. 2i**). This antibody did not block YFV infection of the LRP-KO cells, demonstrating its specificity (**Extended Fig. 4f**).

**Figure 2.**
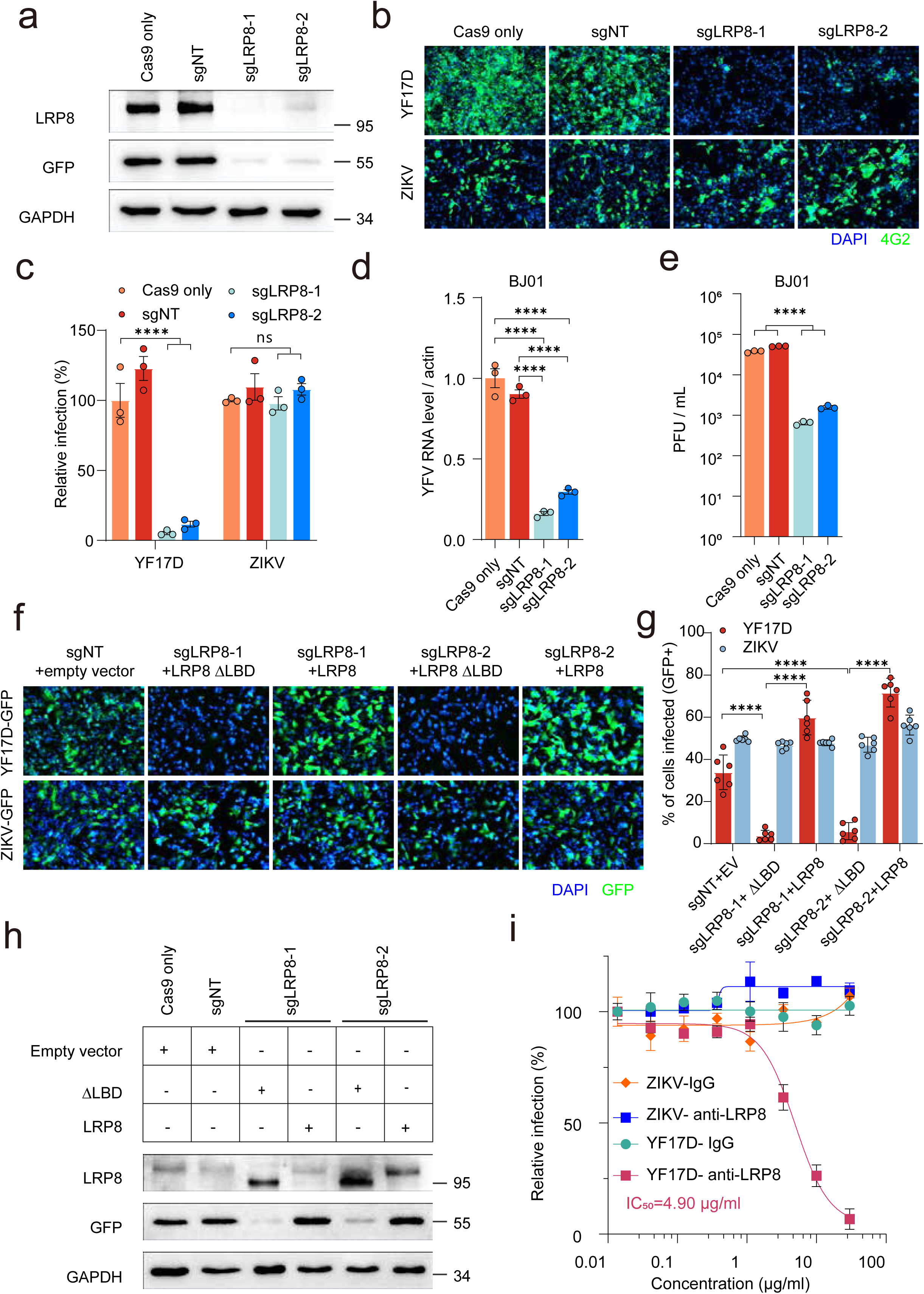
YFV infection requires LRP8 in U87-MG cells. **a**, Cas9/sgRNAs or Cas9 only were expressed in U87MG; single clones were selected. Knockout cells were infected with 17D-GFP for two days and analyzed by Western blot. sgNT: non-targeting sgRNA; sgLRP8-1/2: sgRNAs targeting human LRP8. **b– c**, LRP8 knockout cells or control cells were infected with live 17D or ZIKV for two days, fixed, stained with 4G2 and DAPI, imaged, and quantified. **d-e**, Knockout cells were infected with the clinical BJ01 (0.2moi), viral RNA was quantified by RT–qPCR and viral titer was quantified by focus-forming assay. **f–h**, U87MG LRP8-KO cells were reconstituted with LRP8 WT or LRP8-ΔLBD by electroporation and infected with 17D-GFP or ZIKV-GFP; imaging and quantification as above. **h**, Western blot analysis of LRP8 and GFP as in the 17D-GFP infected group of **f**. **i**, U87MG cells pretreated with an anti-LRP8 antibody or isotype IgG were infected with YF17D or ZIKV and quantified by 4G2 staining. Statistical analysis was performed using one-way analysis of variance (ANOVA) with Dunnett’s multiple comparisons test(d,e), two-way analysis of variance (ANOVA) with Turkey’s multiple comparisons test (c, g). The statistics shown are Mean ± SEM. *: P < 0.05; **: P < 0.01; ***: P < 0.001; ****: P < 0.0001, ns, not significant. The replicate in the figure indicates biological replicates, the experiments were repeated at least three times.

### LRP8 promotes the entry of YFV

We next examined whether LRP8 affects the entry step using 17D. Viral binding to cells was measured after 90 min at 4 °C, while internalization was measured after 2 h at 37 °C. Overexpression of LRP8 significantly boosted binding and subsequent internalization (**Fig. 3a**). Accordingly, LRP8 knockout in U87-MG cells reduced YFV binding and internalization (**Fig. 3b and 3c**). An E-targeting antibody similarly inhibited binding, serving a positive control (**Fig. 3b**). An imaging approach further showed that LRP8 expression promoted binding and internalization of YFV virus-like particles (VLPs). Alexa Fluor 647-labeled YF17D VLPs were incubated with cells at 37 °C for 30 min, washed extensively, and membranes were marked with FITC-WGA. Compared to luciferase or LRP8-ΔLBD, LRP8 significantly increased the number of VLPs at the membrane and in the cytoplasm (**Fig. 3d and 3e**), the effect exceeded that of TIM1. In contrast, VLPs bearing Zika prM/E were not significantly affected by LRP8 (**Fig. 3e and 3f**). Again, the LBD was required: LRP8-ΔLBD and LDLR did not promote YFV entry (**Fig. 3d and 3e**).

**Figure 3.**
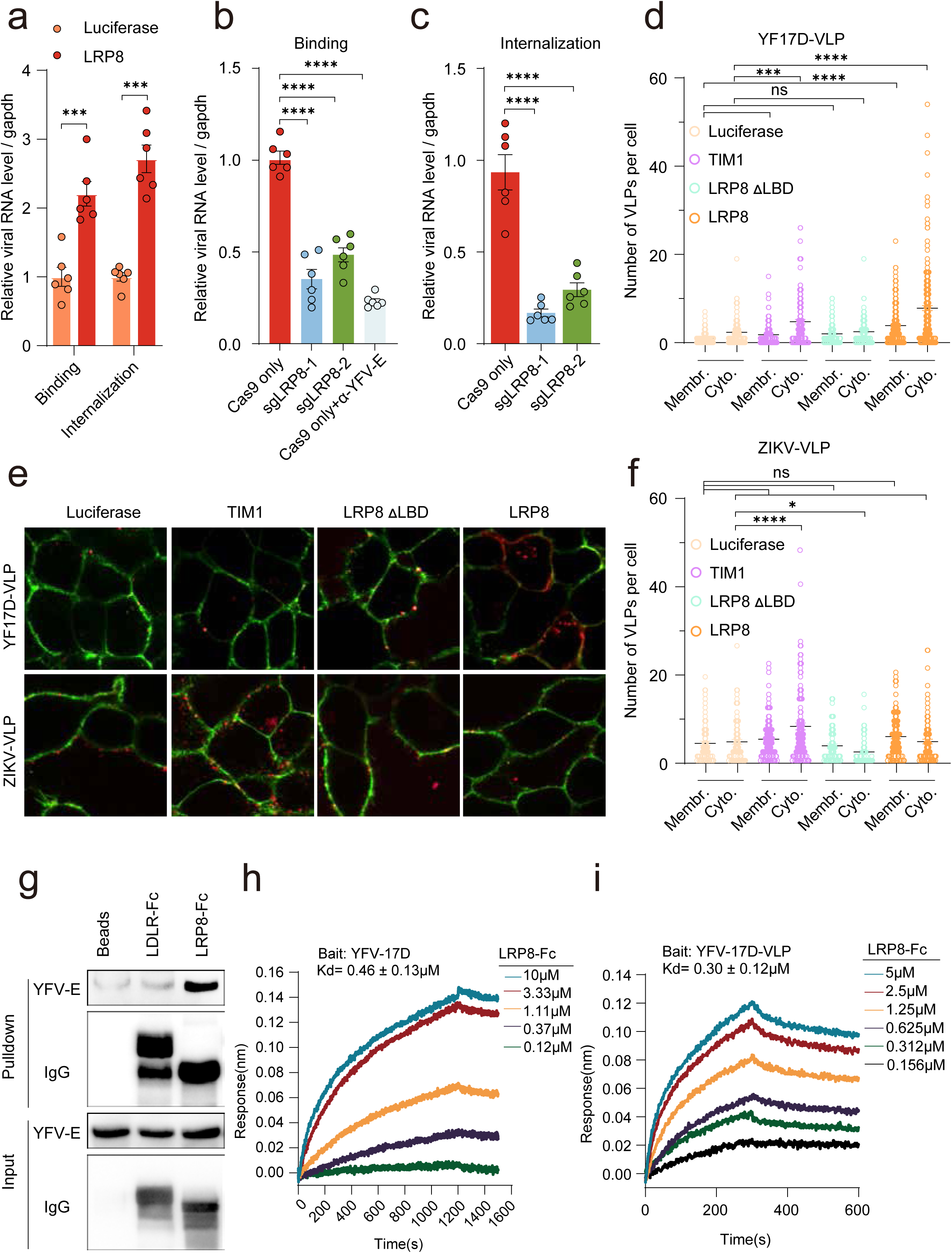
LRP8 promotes YFV entry by binding to its envelope protein. **a**, HEK293T cells stably expressing luciferase or LRP8 were infected with YF17D at MOI 1 either at 4 °C for 45 min to measure binding or at 37 °C for 1 h to measure internalization. **b–c**, U87MG LRP8-KO and Cas9-only control cells were tested similarly; a pan-flavivirus E antibody served as a blocking control. **d–f**, HEK293T cells expressing LRP8, LRP8-ΔLBD, TIM1, or luciferase were treated with Alexa Fluor 647–labeled VLPs bearing YFV 17D E (**d, e**) or ZIKV E (**f**). FITC–WGA was added to label cell membranes before imaging. **g**, Pulldown of YF17D E protein by immobilized LRP8 LBD-Fc. **h, i,** BLI measurement of immobilized live YF17D (**h**) or YF17D VLP (**i**) binding to soluble LRP8-LBD. Statistical analysis was performed using two-tailed unpaired t-tests (a), one-way analysis of variance (ANOVA) with Dunnett’s multiple comparisons test(b, c), two-way analysis of variance (ANOVA) with Turkey’s multiple comparisons test (d, f). The statistics shown are Mean ± SEM. *: P < 0.05; **: P < 0.01; ***: P < 0.001; ****: P < 0.0001, ns, not significant. The replicate in the figure indicates biological replicates, the experiments were repeated at least three times.

### LRP8 interacts with the envelope protein of YFV

We next tested whether LRP8 interacts with the YFV envelope protein, a prerequisite for a bona fide receptor. We purified Fc-tagged LRP8-LBD (LBD-Fc) and performed pulldown assays. YFV envelope protein was specifically pulled down with LRP8 LBD-Fc but not LDLR LBD-Fc (**Fig. 3g**). Biolayer interferometry (BLI) further measured the interaction. LRP8-LBD immobilized on the sensor bound purified YF17D particles with fast, strong kinetics, whereas LDLR-LBD did not (**Extended Fig. 5**). Conversely, immobilized YFV particles bound soluble LRP8-LBD, allowing estimation of an apparent affinity (Kd = 0.46 ± 0.13μM) (**Fig. 3h**). YF17D VLPs containing only E and M proteins also bound LRP8-LBD with similar affinity (Kd = 0.30 ± 0.12μM) (**Fig. 3i**), together with the pull-down experiment, supporting a direct E–LRP8 interaction mediated by the LBD of LRP8.

### Profiling of potential receptor usages by YF17D and clinical YFV strains

It has been demonstrated that VLDLR, another LDLR family protein highly homologous to LRP8, can serve as a receptor for alphaviruses. We therefore tested whether this applies to YFV. While LRP8 strongly promoted 17D, BJ01 and Asibi strains, VLDLR promoted infection by YF17D to a level similar to LRP8 but showed a weaker effect on YFV-BJ01 or Asibi compared with LRP8 (**Fig. 4a, 4b and 4c**). Screening other LDLR family members revealed that, in addition to LRP8 and VLDLR, only LRP4 promoted YF17D infection when overexpressed (**Fig. 4a and Extended Fig. 6**). Similar to VLDLR, LRP4 exhibited a weaker effect on BJ01 (**Fig. 4b**). For LRP1, full-length expression was not observed, likely due to its large size (>500 kDa), precluding conclusions (**Extended Fig. 6**). Interestingly, mouse LRP8 and mouse VLDLR, which promoted YF17D as strongly as human LRP8, supported Asibi pseudovirus and BJ01 only weakly if at all (**Fig.4a and 4b**). Because YF17D was passaged in mouse brain before chick embryos, adaptation to mouse LRP8 likely occurred during passage.

**Figure 4.**
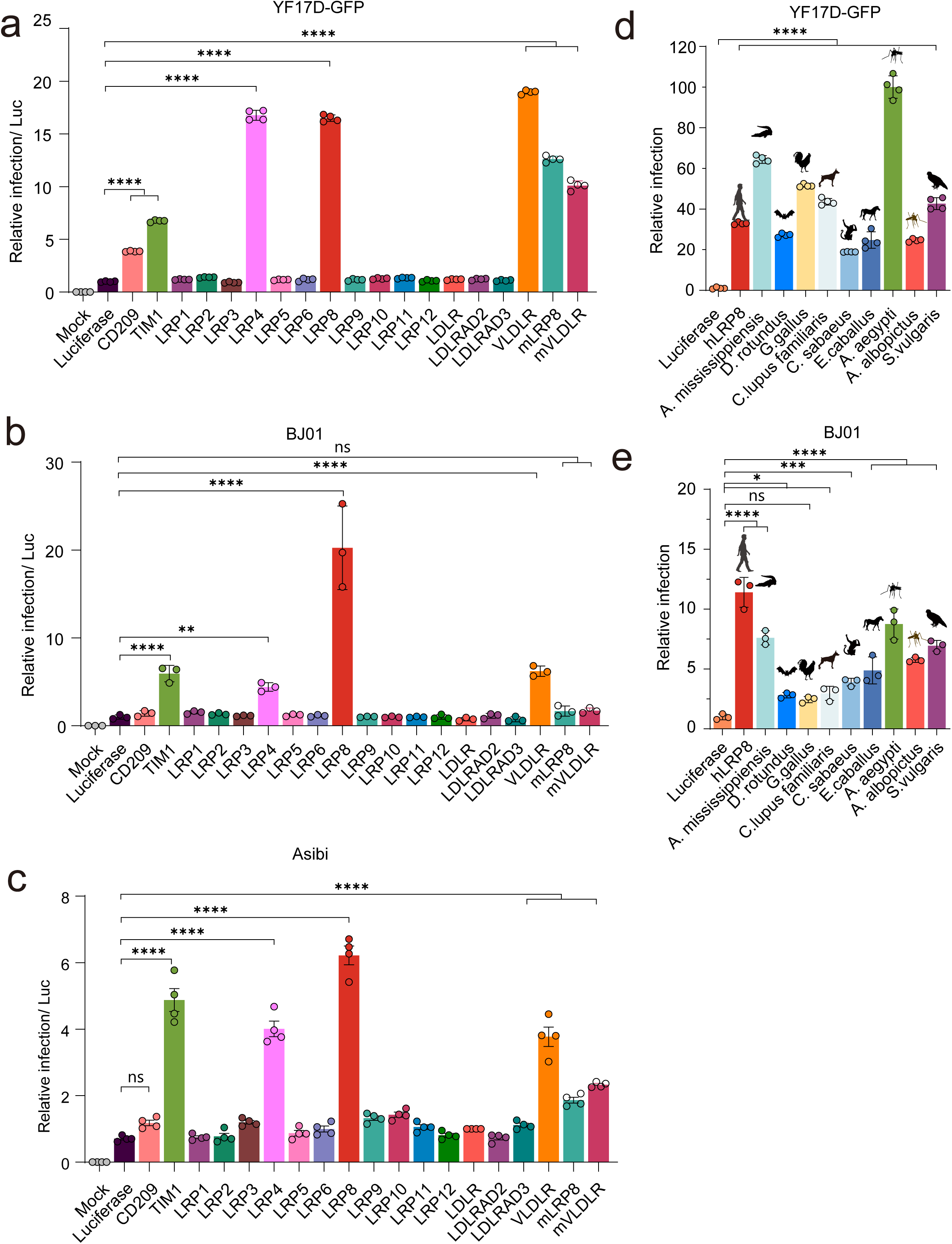
Effects of LDLR family members and LRP8 homologs on YFV infection. **a, b, c**, HEK293T cells transfected with LDLR family members were infected with YF17D-GFP or BJ01 or Asibi for 48 h and analyzed by fluorescence microscopy (YF17D-GFP) or qRT-PCR (BJ01 and Asibi). **d**, **e**, LRP8 homologs from different species were expressed in HEK293T cells and infected with YF17D-GFP or BJ01 for 48 h and analyzed by fluorescence microscopy (YF17D-GFP) or qRT-PCR (BJ01). Statistical analysis was performed using one-way analysis of variance (ANOVA) with Dunnett’s multiple comparisons test (a-e). The statistics shown are Mean ± SEM. *: P < 0.05; **: P < 0.01; ***: P < 0.001; ****: P < 0.0001, ns, not significant. The replicate in the figure indicates biological replicates, the experiments were repeated at least three times.

We also tested a panel of LRP8 homologs from different species by overexpression (**Fig. 4d**, **4e and Extended Fig. 7 and 8**). Most homologs facilitated YFV infection, indicating strong conservation from mammals to mosquitoes and C. elegans. In contrast, BJ01 usage of LRP8 homologs was more restricted, consistent with the shared non-conserved residues that distinguish Asibi from YF17D (**Extended Fig. 9**).

### LRP8 is expressed in human liver and mediates YFV infection in primary human hepatocytes and in a mouse model

LRP8 is well known to be highly expressed in brain, but its expression in human liver is less well characterized. Because liver is a key target organ of YFV, we assessed hepatic expression of LRP8 and homologs. Immunohistochemistry with two different LRP8 antibodies of human liver samples, showed clear LRP8 expression at levels comparable to brain (**Fig. 5a and Extended Fig. 10**). This was corroborated by Western blots and single cell sequencing data of human liver samples (**Fig. 5b and 5c)**. In contrast, LRP4 and VLDLR were low (**Fig. 5b and 5c)**.

**Figure 5.**
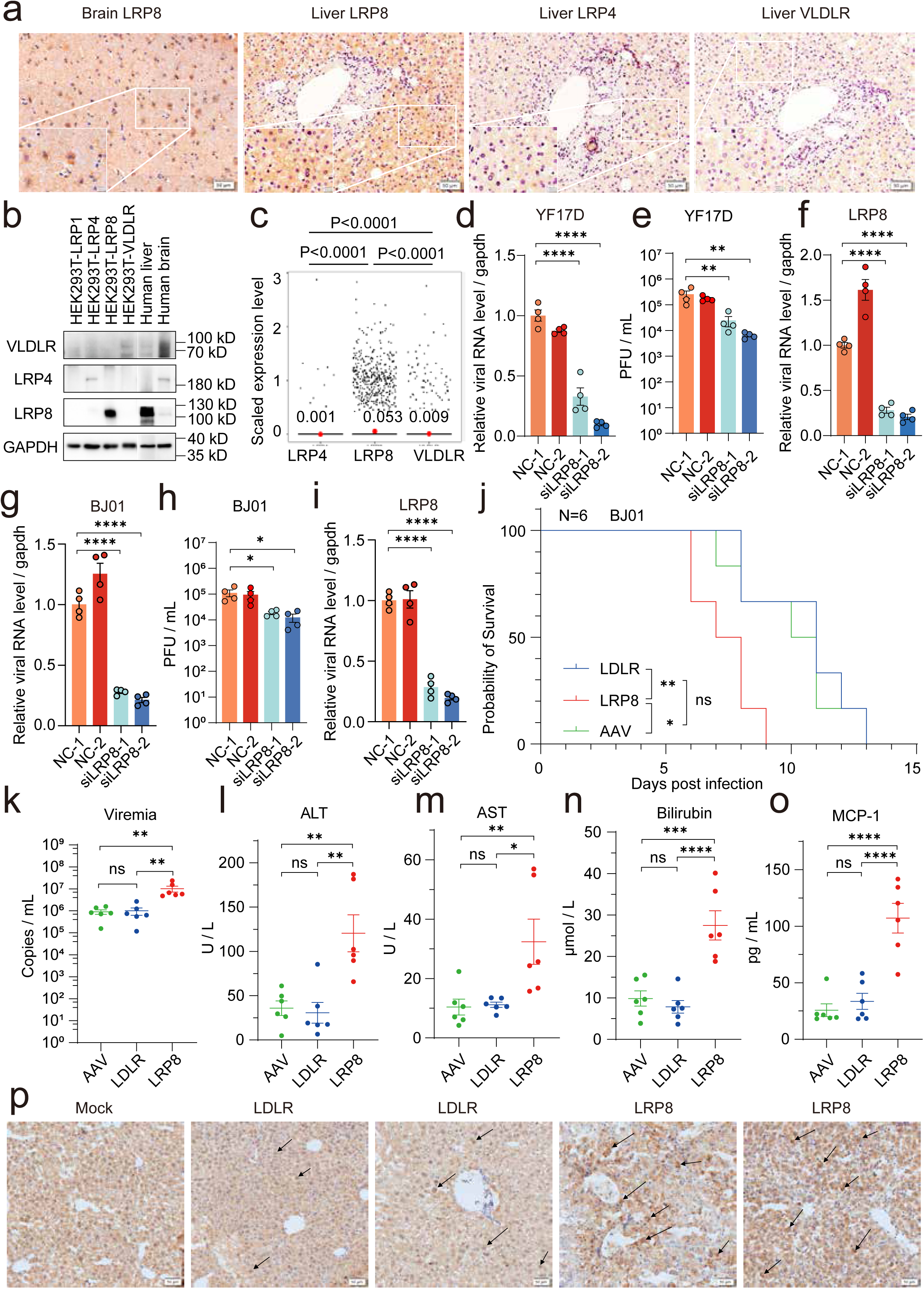
LRP8 is expressed in human liver and mediates YFV infection in primary human hepatocytes and in a mouse model. **a**, LRP8, LRP4 and VLDLR expression were determined in human brain or liver by immunohistochemistry staining with corresponding antibodies. **b**, Scaled expression levels of LRP8, LRP4 and VLDLR in human liver single cell RNAseq data. The red dot within each column represents the mean expression value across all cells. **c**, Western blots of human liver samples, including crude liver lysates, showed clear LRP8 expression at levels comparable to brain. HEK293T cells overexpressing one of the three genes were included as controls showing the specificity of the antibodies. **d-i**, Primary human hepatocytes were transfected with siRNAs targeting LRP8 or non-targeting control siRNAs (NC-1 and NC-2) for 48 hours and infected with live YF17D (**d-f**) or BJ01 (**g, i**) for 48 hour, and the viral RNA and LRP8 mRNA level from cell total RNA were quantified via qRT-PCR (**d, f, g, j**). The virus titers of the supernatant were quantified by plaque assay (**e, h**). **j-p**, In AG129 mice, 2x 10^11^ genome-copies of AAV-TBG expressing either LRP8 or LDLR and an empty AAV were injected by tail vein into each mouse. Three weeks later, YFV BJ01 was injected intraperitoneally. The survival curve of the mice were shown in j. Five days post YFV injection, the blood was drawn from the mice to measure viral load (**k**) and levels of ALT and AST enzymatic activities and levels of bilirubin and MCP-1 (**i-o**). The livers of a mock mouse and two mice each from the LDLR and LRP8 group died at day 7 (LRP8 group) or day 10 (LDLR group) were processed for immunohistochemistry imaging with DAB staining of YFV E protein with an anti-E antibody, asterisks indicate sites of positive staining (**p**). Statistical analysis was performed using one-way analysis of variance (ANOVA) with Dunnett’s multiple comparisons test (b, d-i, k-o),log-rank (Mantal Cox) test (j). The statistics shown are Mean ± SEM. *: P < 0.05; **: P < 0.01; ***: P < 0.001; ****: P < 0.0001, ns, not significant. The replicate in the figure indicates biological replicates, the experiments were repeated at least twice.

Given LRP8 expression in the liver, we tested LRP8 knockdown in primary human hepatocytes using siRNAs. Two siRNAs significantly reduced infection of PHHs by YFV-17D and YFV-BJ01 relative to controls as measured by both cellular viral RNA levels and viral titers of the media, supporting a key role for LRP8 in human liver (**Fig. 5d-5i**). Accordingly, LRP8 decoy protein also blocked YFV infection of PHHs (**Extended Fig. 11a**).

To test human LRP8 in vivo, we used Adeno-associated virus (AAV) to overexpress LRP8 in the liver of AG129 mice, which are deficient in the interferon signaling pathway and thus useful for modeling YFV-associated viscerotropic disease^8^. The overexpression levels were confirmed by Western blots of the mouse livers (**Extended Fig. 11b**). Compared with LDLR overexpression (which does not promote YFV entry), LRP8 significantly aggravated pathology after BJ01 infection (**Fig. 5j**). Survival of mice with liver-specific LRP8 expression was significantly worse than with LDLR, correlating with higher viral titers, elevated MCP-1, and increased ALT/AST/bilirubin (**Fig. 5k-5p**). Survival in the LDLR group was similar to empty-AAV controls, further supporting LRP8-mediated enhancement in vivo.

### LRP8 mediates YFV 17D infection in mosquitoes

Because mosquitoes have a single LRP8/VLDLR-like gene (**Extended Fig. 7**), we used in vivo gene knockdown via injection of synthesized double-stranded RNA (dsRNA) targeting *Aedes aegypti* VLDLR in the thorax. Four dsRNAs effectively knocked down VLDLR, and correspondingly blocked infection by YF17D as measured by RT-qPCR (**Fig. 6a and 6b**). Consistent with cellular experiments, this block was specific to YFV and had no effect on Zika virus infection (**Fig. 6c and 6d**). These results, together with the overexpression data, confirm that the mosquito LRP8 homolog is a key factor mediating YFV infection in a natural host.

**Figure 6.**
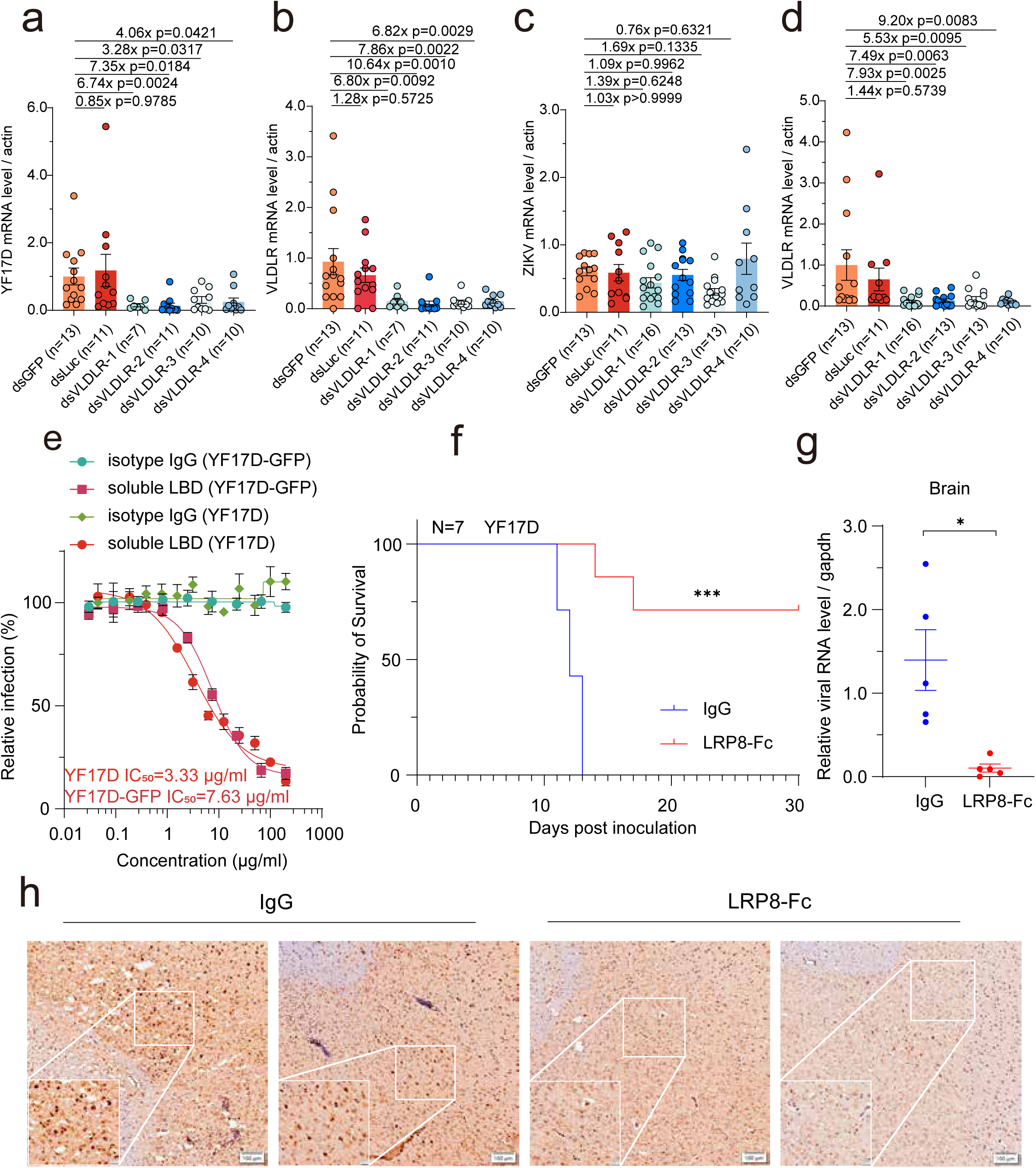
LRP8 function in YFV infection in animal models. **a–b**, *Aedes aegypti* mosquitoes injected with dsRNA targeting VLDLR or control dsRNA for three days were then infected with YF17D. Four days post infection, viral RNA (**a**) and VLDLR mRNA (**b**) were quantified by qRT–qPCR. **c–d**, dsRNA-treated mosquitoes were infected with ZIKV. **e**, YF17D or 17D-GFP was treated with soluble LRP8 LBD-Fc or isotype IgG and used to infect U87MG cells; infection was quantified by immunofluorescence or GFP. **f**, C57BL/6 mice were injected intracerebrally with YF17D preincubated with LRP8 LBD-Fc or isotype IgG; survival was recorded daily. **g**. Brain from mice of **f** was drawn to measure viral load by qRT-PCR. **h**. In a repeat experiment of **f**. mice at day 10 post infection were sacrificed and the brain of the mice were processed for immunohistochemistry imaging with DAB staining of YFV E protein with an anti-E antibody. Statistical analysis was performed using one-way analysis of variance (ANOVA) with Dunnett’s multiple comparisons test (a-d), log-rank (Mantal Cox) test (f), two-tailed unpaired t-test with Welch’s correction (g). The statistics shown are Mean ± SEM. *: P < 0.05; **: P < 0.01; ***: P < 0.001; ****: P < 0.0001, ns, not significant. The replicate in the figure indicates biological replicates, the experiments were repeated at least twice.

### Purified Ectodomain of LRP8 protein blocks YFV infection

YFV infection of immunocompetent mice causes mortality when injected intracerebrally, a well-established model for assessing virulence. Using this model, we employed a decoy approach to test LRP8 function in mice. Purified Fc-tagged LRP8-LBD demonstrated dose-dependent, specific inhibition of YFV infection in U87-MG cells (**Fig. 6e**). C57BL/6 mice were injected intracerebrally with 17D preincubated with either purified Fc-tagged LRP8-LBD or an isotype IgG control. Mortality was significantly decreased in the LRP8-LBD group compared with controls (**Fig. 6f**), so are the viral loads shown by qRT-PCR and immunohistochemistry staining (**Fig. 6g and 6h**), confirming that soluble LRP8-LBD can block 17D infection and supporting its role as a receptor that directly binds YFV E on the virion surface.

## Discussion

Virus receptors provide essential footholds for host-cell entry. Discovering them enables delineation of viral life cycles, interpretation of pathogenesis, and development of therapies. LRP8 as a YFV host factor is conserved from mosquitoes and ticks to non-human primates and humans. This suggests an advantage to a one-receptor-for-many-hosts strategy, with fewer evolutionary constraints. It is known that YFV 17D can infect the midgut of Aedes aegypti mosquitoes but does not disseminate to the salivary glands and therefore cannot be transmitted, suggesting a complex mechanism behind tissue tropism and the possibility of other receptors involved in this vector^29,30^. Closest relatives of YFV such as Sepik and Wesselsbron viruses do not appear to engage LRP8. Recent work also suggests that tick-borne encephalitis virus (TBEV) uses LRP8^31^, and phylogenetic analysis indicates that LDLR-family usage has arisen in distant virus families, likely via independent evolution (**Extended Fig. 12**). Often multiple LDLR-family members can support entry for a given virus. For YFV, our data indicate that, in addition to LRP8, LRP4 and VLDLR can promote entry, possibly in different cells or tissues. This receptor redundancy highlights the advantage of overexpression screens, which can overcome redundancy that confounds loss-of-function approaches. That the soluble LRP8 LBD can block YFV infection, together with prior studies, raises the possibility of LRP8-based protein therapeutics or prophylactics for multiple arboviruses. Finally, wild-type YFV strains such as Asibi or BJ01 do not respond to mouse LRP8 or VLDLR as robustly as 17D, consistent with mice not being the natural host and suggesting adaptation to mouse LRP8 during historical passaging to generate 17D. These findings illuminate the evolutionary path underlying one of the most successful vaccines and may inform prevention strategies for other flaviviruses. Our data showing that expression of human LRP8 in mouse livers can aggravate the pathogenesis may support an improved mouse model of YFV infection with more consistent visceral phenotype than the current mouse model^32^.

## Acknowledgements

This work was supported by a grant from the State Key Research Development Program of China to X.T. (2021YFA1302503 and 2022YFE0102200). We thank Lin Mei and Bin Luo for reagents, Wang L, Ding N and Yan X for technical assistance, Zhang Yuhang and Zhang Guigen for editing the manuscript.

## Author contributions

X.T. conceived and supervised the project. M. M., Y. Yang, Z. Zhang, Y. Yin, J. T., J. C., Y. G., Z. W., D. W., Y. J., Y.C., H. W. conducted the experiments. Zhiyuan Zhang, L. L. and Y. P. contributed computational analysis, R. L., Q. H., Z.D., Q. H., C. S., Y. Z J. L., W. T., L. M. and G. C. provided reagents, guidance and facility support. M.M. and X.T. wrote the manuscript with input from all other authors.

## Competing interests

The authors declare no competing interests.

## Materials and Methods

### Cells and viruses

We maintained HEK293T (human kidney epithelial, ATCC CRL-11268), HEK293FT (human kidney epithelial, Thermo Fisher Scientific, R70007), U87MG (human glioblastoma, ATCC HTB-14), Vero (Cercopithecus aethiops kidney, ATCC CCL-81), A549 (human lung epithelial, ATCC CCL-185), Huh7 (human hepatoma, CRL-10741), K562 (chronic myelogenous leukemia cell line, CCL-243), C6/36 (Aedes albopictus larva, ATCC CRL-1660), and BHK-21 (ATCC CCL-10) in DMEM (Gibco) supplemented with 10% fetal bovine serum (FBS), Primary human hepatocytes (Shenzhen Liver Biotechnology Co., Ltd.), 1% GlutaMAX (Thermo Fisher Scientific, 35050061), and 1% penicillin– streptomycin (Thermo Fisher Scientific, 151401148). We confirmed the absence of mycoplasma by monthly testing using myco-Lumi (Beyotime, C202985S) after ZellShield treatment (Minerva Biolabs, 13-0050).

Wild-type live flaviviruses used were YFV (clinical strain BJ01、Asibi and the live-attenuated vaccine strain YF17D) and ZIKV (strain SZ01). Viruses were propagated in C6/36 or Vero cells and titrated in BHK-21 by plaque assay.

### Antibodies

Reagents included: anti-YFV Env (Creative Diagnostics, CABT-YN1194), anti-ZIKV Env (Creative Diagnostics, CABT-YN1207), LRP4 (Invitrogen, MA5-45577), LRP4 (GeneTex, GTX637988), ApoER2/LRP8 (Santa Cruz, sc-293472), Anti-VLDLR (GeneTex, GTX79552), LRP8 (Thermo Fisher, MA5-38566), ApoER2/LRP8 (Abclonal, A10517), pan-flavivirus Env (Creative Diagnostics, CABT-YN1131), 4G2 (Millipore, MAB10216), anti-Tim1 (Abclonal, A2831 and A26862), HRP-conjugated anti-rabbit (ZSGB-Bio, ZB-2301) anti-mouse (ZB-2305), and anti-human (W4038), Alexa Fluor 488 goat anti-mouse (A-11001), and Alexa Fluor 568 goat anti-rabbit (A-11011).

### Reporter pseudovirus particle generation

To generate replication-incompetent reporter pseudovirus particles (RPPs), we used a DNA-based WNV replicon lacking the C-prM-E coding region (pWNVrep dCprME-GFP) co-transfected with a plasmid expressing flaviviral CprME. We cloned CprME of ZIKV strain SZ01 (GenBank KU866423), YF17D (GenBank OR509732), Asibi (GenBank MF405338), CNYF01 (GenBank KU921608),

WESSV (GenBank NC_012735), SEPV (GenBank DQ837642), Dengue virus type 2 (GenBank NC_001474), WNV (GenBank NC_001563.2), JEV (GenBank NC_001437.1), and mutants into pcDNA3.1 under a CMV promoter. HEK293T cells (5×10^6 per 15-cm dish) were seeded 12–18 h before transfection. Standard medium (DMEM + 10% FBS, penicillin–streptomycin, GlutaMAX) was used; cells were 50–60% confluent at transfection. We co-transfected 30 μ g pWNVrep dCprME-GFP and 10 μ g pcDNA3.1-CprME using Neofect reagent, replaced medium at 8–12 h, and collected supernatants 3 days post-transfection. Supernatants were centrifuged (4,000 rpm, 5 min), filtered (0.45 μm), and stored at −80 °C.

For purification, supernatants at 48 and 72 h were clarified (4,000 rpm, 10 min) and concentrated by Amicon Ultra-30 kDa units or purified on 10–60% continuous sucrose gradients (200,000 g, 2 h). Bands were collected and washed three times with ice-cold PBS using Amicon-30 kDa filters. Preparations were stored at −80 °C (long-term) or 4 °C (≤2 weeks). Antigen content was quantified by YFV ELISA (COIBIO, CB11927-Hu).

### Production and Purification of YF17D Live-Attenuated Virus

YF17D was propagated in C6/36 at MOI 0.5 and 28 °C. Six days post-infection, supernatant was harvested and centrifuged at 4,000 rpm for 30 min to remove debris, filtered (0.45 μm), and precipitated with 40% PEG8000 at 4 °C overnight. Pellets were collected (11,000 rpm, 30 min), resuspended in TNE buffer (Shyuanye R20992), and purified by ultracentrifugation on a 24% sucrose cushion (105,000 g, 2 h, 4 °C). Further purification used a 10–35% potassium tartrate gradient (32,000 rpm, 2 h, 4 °C). Visible bands were collected and concentrated by ultrafiltration (100 kDa MWCO).

### VLP labelling and imaging

Purified YF17D or ZIKV VLPs were labelled with Alexa Fluor 647 NHS ester in 0.1 M sodium bicarbonate (pH 8.3) for 30 min at room temperature in the dark. Reactions were quenched with 50 mM Tris–HCl (pH 7.5), and unbound dye was removed by centrifugal filtration (100 kDa MWCO). Input was normalized by envelope antigen content. HEK293T cells stably expressing luciferase, TIM1, LRP8-ΔLBD, or LRP8 were used. Binding assays were performed at 4 °C for 45 min with six PBS/2% FBS washes; internalization was at 37 °C for 30 min with optional proteinase K treatment (500 ng/mL, 15 min, 37 °C). Cells were stained with FITC–WGA (5 μg/mL, 10 min, 4 °C) and DAPI, fixed in 4% paraformaldehyde, and imaged by confocal microscopy. Single-cell VLP counts were quantified using CellProfiler.

### Bio-layer interferometry analysis

BLI analysis were performed at 25°C using an Octet R8 biosensor system (Sartorius). YFV E antibody (GTX638626, GeneTex) was mixed and incubated with biotin (#G-MM-IGT, Genemore) at a molar ratio of three biotins to one protein at room temperature for 30min. Excess biotin was removed using a desalting column(#G-MM-IGT, Genemore). Biotinylated protein (25μg·mL−1) was immobilized onto the SA Biosensors (Sartorius #18-5019) for 3 min and then used to capture YFV VLPs or YFV-17D viruses. The tips were washed with the wash buffer (20 mM HEPES, 150 mM NaCl, pH 8.0) for 3 min to obtain a baseline reading, then the biosensors were dipped into wells containing the LRP8-Fc at various different concentrations for the indicated time. BLI data analysis was performed by using the software Octet (version 12.2, Sartorius) in the standard 1:1 binding mode. Raw binding data were exported and plotted in GraphPad Prism8.0.

### Genome-wide ORF overexpression screen

The pooled human ORF library contains 20,432 clonal ORFs mapping to 13,833 genes. HEK293T cells (1×10^7 per 150 mm dish) were transfected with lentiviral ORF sublibraries and packaging plasmids (SV40 VSVg, Gag/Pol, TAT, Rev; 1:1:1:1) using Neofect. Supernatants were collected at 36–48 h, aliquoted, and stored at −80 °C. Sub-library titers were determined by puromycin selection. Five million HEK293T cells were transduced at MOI 0.2, selected with puromycin (2 μg/mL, 3 days), mixed in equal numbers, and infected with YF17D-GFP at MOI 0.2 or mock-infected. At 24 hpi, GFP+ and GFP− cells were sorted by FACS. Genomic DNA was extracted, barcodes amplified, and sequenced (HiSeq-PE150). Reads were trimmed with Cutadapt, aligned with Bowtie2, processed with SAMtools, counted with a custom Python script, normalized with DESeq2, and log2 fold changes calculated relative to GFP− samples. Scores integrated read counts and fold change (see Supplementary Data Set).

### Viral entry assay

For binding assays, HEK293T-Luc, HEK-LRP8iso3, U87MG-Cas9 only, U87MG-sgNT, U87MG-sgLRP8-1, and U87MG-sgLRP8-2 cells (0.1 M per well, 24-well plates) were incubated with YF17D, BI01 and / or Asibi at MOI 1 at 4 °C for 45 min and washed six times with ice-cold PBS/2% FBS. An anti-YFV Env antibody (Creative Diagnostics, CABT-YN1194) served as a blocking control (virus preincubated with 30× diluted antibody at 37 °C for 1 h). Cells were lysed in RLT buffer (R4111, Magen) and RNA extracted for RT–qPCR. For internalization, after binding and washing, prewarmed medium (37 °C, 2% FBS, 15 mM NH4Cl) was added and cells were incubated for 1 h at 37 °C. Residual surface virions were removed with proteinase K (500 ng/mL, 15 min, 37 °C) and ice-cold PBS/2% FBS before RNA extraction.

### Ectopic overexpression experiments

For stable or transient expression assay, ORFs of LRP8 (GenBank NM_017522.5), and LDL family: LRP3 (GenBank NM_002335.4), LRP4 (GenBank NM_002334.4), LRP5 (GenBank NM_002335.4), LRP6 (GenBank NM_002336.3), LRP9 (GenBank NM_003105.6), LRP10 (GenBank NM_014045.5), LRP11 (GenBank NM_032832.6), LRP12 (GenBank NM_013437.5), LDLR (GenBank NM_000527.5), LDLRAD2 (GenBank NM_001013693.3), LDLRAD3 (GenBank NM_174902.4), VLDLR (GenBank NM_013703), CD209 (GenBank NM_021155.4), TIM1 (GenBank NM_012206.3) were cloned into a CMV-driven lentivector with FLAG/HA tags. Codon-optimized LRP1 (GenBank NM_002332), LRP2 (GenBank NM_004525.3), *A. aegypti* LRP8 (GenBank XP_021711662), *Aedes albopictus* VLDLR (GenBank XM_029869759.2), *Equus caballus* LRP8 (GenBank XP_023485552), *Sturnus vulgaris* LRP8 (GenBank XM_014870608.1), *Canis lupus familiaris* LRP8 (GenBank XM_038666202.1), *D. rodundus* bat LRP8 (GenBank XM_053920761.1), *Alligator mississippiensis* LRP8 (GenBank XM_014598721.3), *Gallus chicken* LRP8 (GenBank NM_205186.3), *Ixodes scapularis* VLDLR (GenBank XM_029983583.4), mouse LRP8 (GenBank NM_001369051.1), mouse VLDLR (GenBank NM_013703.2), were synthesized into pcDNA3.1 with C-terminal 3×HA. Lentiviral transduction and puromycin selection (2 μg/mL, 3 days) generated stable lines. For HA-tagged constructs, surface expression was confirmed by FACS using surface staining. The target protein sequences from NCBI were downloaded and performed multiple sequence alignment by Clustal W using the Geneious software package.

### Genetic knockdown and knockout validation

Specific siRNAs targeting LRP8 were synthesized by Jintuosi. In PHH cells (Liver Biotechnology), 100nM siRNAs were transfected using RNAiMax. After 48 hours, the medium was replaced with PHH maintenance medium, and the cells were infected with YF17D and BJ01. Another 48 hours later, the cells were collected to detect viral RNA levels and the knockdown efficiency of LRP8 mRNA. The siRNA sequences are as follows:

siLRP8-1: CCUUGAAGAUGAUGGACUATT,
siLRP8-2: CTGGACTGACTCGGGCAATAA.

The single guide RNAs targeting LRP8 (Genebank NG011517) were designed as follows,

F1, caccgCGTCACACTTCCACCGTTCG,
R1, aaacCGAACGGTGGAAGTGTGACGc;
F2: caccgACTGTCTGCACAGGTCTTCT,
R2: aaacAGAAGACCTGTGCAGACAGTc).

Then these guide RNAs were cloned into lenti-CRISPRv2 plasmid (Addgen#52961). Lentivirus was collected from the supernatant of HEK293T cells transfected with lenti-CRISPRv2 in addition of packaging plasmids aforementioned. LRP8 knockout (KO) clones were generated by single guide RNA in U87MG and selected with 2 μg/mL puromycin for 5-7 days. Further sgLRP8 cells were pooled in 96-well plates at a density of 0.5 cell per well to obtain single-cell clones. Each clone was validated by Western blot or Sanger sequencing.

### Pull-down assay

Purified soluble LRP8-Fc (10 μg) or isotype IgG was incubated with 30 μL protein A/G magnetic beads (Thermo Fisher Scientific) in PBS/0.1% Tween-20 at 4 °C for 6 h. After removing unbound proteins, beads were incubated overnight at 4 °C with clarified lysates from YF17D-infected Vero cells, washed eight times with PBS/0.1% Tween-20, and eluted by boiling in SDS-PAGE buffer for immunoblotting.

### Protein purification

LRP8 LA1237 DNA was inserted into a vector encoding a C-terminal human IgG Fc fusion and transfected into HEK293F (3.0×10^6 cells/mL). Supernatants (60h after transfection) were purified on Protein A Sepharose 4B, washed with 20 mM HEPES/150 mM NaCl (pH 8.0), and eluted with 20 mM HEPES/150 mM NaCl/0.1 M glycine (pH 2.8), followed by SEC (Superdex 200 10/300 GL) in 20 mM HEPES/150 mM NaCl (pH 8.0). LDLR-Fc was purified similarly. For VLDLR LA1-8, co-transfection with RAP enabled on-column Ca2+-dependent refolding after extensive EDTA washes.

### Decoy protein and antibody-mediated blocking assay

Purified LRP8-Fc (200 μg/μL stock) or isotype IgG was serially diluted, preincubated with YF17D or ZIKV (pseudovirus or live) at 37 °C for 1 h, then added to cells. For pseudovirus, GFP+ cells were quantified at 24 h by high-content imaging. For live virus, cells were fixed at 48 h and stained with 4G2 and fluorescent secondary antibodies. For antibody blocking, anti-LRP8 or isotype controls were incubated with cells for 1 h at 37 °C before infection.

### Immunoblotting

Cells and tissue samples were lysed in 1% NP-40 buffer on ice. Protein concentrations were determined by BCA, samples denatured in 5× SDS-PAGE buffer (10 min, 95 °C), resolved by SDS-PAGE, transferred to nitrocellulose, blocked in TBST/5% milk, incubated with primary antibody overnight (4 °C), washed, and probed with HRP-secondary antibodies for 1 h before detection.

### Immunofluorescence

Eight thousand HEK293T or U87MG cells per well in 96-well plates were infected with YF17D (MOI 0.3) or ZIKV SZ01 (MOI 0.2) for 24–48 h, fixed in 4% paraformaldehyde, permeabilized with 0.2% Triton X-100 (Promega), and stained with 4G2 (1:1000) and Alexa Fluor 488 secondary (Invitrogen, R37120, 1: 1000). Nuclei were counterstained with DAPI (Invitrogen, 62247, 1:10,000). Images were acquired on a high-content microscope (Cytation 1) and infection ratios quantified.

### Flow cytometry

For membrane protein expression, cells were detached with 10 μM EDTA, stained with anti-LRP8、anti-TIM1 or anti-HA for 30 min at room temperature, washed, stained with AF-488 secondary for 1 h, fixed in 2% paraformaldehyde, washed, and analyzed by flow cytometry. GFP+ cell ratios were quantified after trypsinization for reporter assays.

### Quantitative real-time PCR

DNA/RNA were extracted (Magen, R5111). Two micrograms of total RNA were reverse-transcribed (ABM, G492). qPCR was performed with 2× ChamQ SYBR reagents (Vazyme, Q321) on a Bio-Rad CFX96. qRT-PCR data were normalized to GAPDH. The primer sets were as follows:

*GAPDH*-qF: CGGAGTCAACGGATTTGGTCGTAT;
*GAPDH*-qF: AGCCTTCTCCATGGTGGTGAAGAC;
*YF17D*-qF: ATGGCAGGAGGATTGTGGTG;
*YF17D*-qR: GTTGTGCGTCCTTGTGGAAC;
YFV BJ01-qF: GCTAATTGAGGTGYATTGGTCTGC;
YFV-BJ01-qR: CTGCTAATCGCTCAAMGAACG;
*YFV-Asibi-*qF*: GCACGGATGTAACAGACTGAAGA;*
*YFV-Asibi-*qR*: CCAGGCCGAACCTGTCAT;*
*LRP8*-qF: CCCATCCCTAATCTTCACCAAC;
*LRP8*-qR: CTAGTGCCACGACATTCTTGAG.

### In vivo mosquito experiments

Four dsRNAs targeting the common region of Aedes aegypti VLDLR (LOC5576021) and controls (luciferase, GFP) were designed (SnapDragon), synthesized by in vitro transcription (MEGAscript, Ambion), and purified. Female mosquitoes were chilled on ice and injected (1 μg/300 nl) into the thorax. Gene silencing was verified by qRT-PCR. For infection, 1000 MID50 of YF17D or ZIKV was microinjected. Mosquitoes were collected at 4 dpi for RT-qPCR.

Six pairs primers were synthysized as follows:

DS1 F: TAATACGACTCACTATAGGGATCCCCGTGATCATACCAAA
DS1 R: TAATACGACTCACTATAGGGCCAGTCCGTCCAGTACATCC
DS2 F: TAATACGACTCACTATAGGGAAACCTGCAGATCGGATGAG
DS2 R: TAATACGACTCACTATAGGGTTTTGCACTGGAACTGATCG
DS3 F: TAATACGACTCACTATAGGGCCTGCAGATCGGATGAGTTT
DS3 R: TAATACGACTCACTATAGGGGCAGGTTTTGTCCTTTTTGC
DS4 F:TAATACGACTCACTATAGGGGTATGCCGGGTTACCTGAGA
DS4 R: TAATACGACTCACTATAGGGTCAGAATCTTGCCCATGTTG
Ds-Luc-F: TAATACGACTCACTATAGGGGGACTTGGACACCGGTAAGA
Ds-Luc-R: TAATACGACTCACTATAGGGGTCCACGAACACAACACCAC
DS-GFP-F:TAATACGACTCACTATAGGGGTGAGCAAGGGCGAGGAG
DS-GFP-R: TAATACGACTCACTATAGGGCATGATATAGACGTTGTGGCTGTT.

### Mouse infection experiment

Five-week-old C57BL/6 mice were anesthetized and secured in a stereotaxic apparatus. For in vivo neutralization, YF17D was premixed with purified soluble LRP8-LBD-Fc or isotype IgG at 37℃ for 1 hours, each group contain six equal gender mice. Ten microliter (200 PFU) was injected into the lateral ventricle (AP +0.2 mm, ML ±1.0 mm, DV −2.0 mm) at 10–20 nL/s. The needle was left in place for 5–10 min to minimize backflow, and incisions were closed. Meloxicam (4 mg/kg, s.c.) was administered for three days. Survival was monitored daily. For viral quantification in brain infection and blood, another independant assay was performed, and all the mice were sacrificed at 10 days post infection, the blood and brain were collected.

In the AAV mouse model, 21 three-week-old AG129 mice were evenly divided into three groups, with equal numbers of males and females. Each mouse received a tail vein injection of 2×10^11^ gc of AAV2/8-TBG-LRP8, AAV2/8-TBG-LDLR, or an AAV2/8-TBG empty vector, packaged by CIBR Vector Core. Three weeks after AAV delivery, one mouse from each group was sacrificed to examine receptor expression in the liver. After confirming successful expression, each mouse was intraperitoneally injected with 10,000 PFU of BJ01, then weighed daily and monitored for health status. On the fifth day after YFV infection, blood was collected via the orbital vein to assess viremia, liver function markers, and inflammatory factors. When the mice were close to death, liver and brain tissues were collected to evaluate tissue viral titers and pathological changes. All animal procedures were Tsinghua or Shenzhen Third Hospital IACUC-approved and complied with institutional and national guidelines.

### Sera biochemistry analysis & ELISA analysis

Alanine transaminase (ALT), Aspartate aminotransferase (AST) and Total Bilirubin (TBIL) in mouse serum was detected using biochemistry assay kits according to the corresponding instructions. All of the reagent kits were manufactured by Nanjing Jian cheng Biotechnology Co., Ltd. The levels of and MCP-1 in mouse serum were measured using commercial ELISA kits manufactured by Sino Biological Biotechnology Co., Ltd. ALT detection kit (C009-2-1, Nanjing Jiancheng Bioengineering Institute Co., Ltd), AST detection kit (C010-2-1, Nanjing Jiancheng Bioengineering Institute Co., Ltd), TBIL detection kit (C019-1-1, Nanjing Jiancheng Bioengineering Institute Co., Ltd), MCP-1 ELISA kit (KIT50368, Sino Biological Biotechnology Co., Ltd).

### Immunohistochemistry

After mouse sacrificed, the tissues were fixed in 4% paraformaldehyde overnight at 4°C, then followed by 20%, 30% sucrose gradient dehydrated 1 day, embedded in OCT (SAKURA, 4583) and cut into 10 μm sections. For Immunohistochemistry (IHC), antigen retrieval was performed by boiling in 0.01 M citrate buffer (pH 6.0) for 10 min. Endogenous hydrogen peroxidase activity was blocked by 3% hydrogen peroxide in PBS for 30 min, followed by incubating in primary antibodies overnight at 4 ℃, HRP-conjugated anti-rabbit secondary antibody at RT (room temperature). DAB staining followed instructions from the DAB detection kit (ZSGB-Bio, ZLI-9019) for 5–10 min at RT, finally counter staining was performed using hematoxylin solution (Solarbio, H8070). Primary antibodies and dilution factors are listed below: anti-YFV Env (Creative Diagnostics, CABT-YN1194, 1:100), anti-LRP4 (GeneTex, GTX637988, 1:100), anti-VLDLR (Abcam, ab302917, 1:100), LRP8 (Thermo Fisher, MA5-38566, 1:100), anti-LRP8 (Abclonal, A10517, 1:50).

### Plaque assay for viral titer identification

Supernatant titration (in PFU mL−1) of each HEK293T over-expressing cells was obtained by plaque assay to determine the amount of infectious viral particles (PFU). The virus titration was performed in BHK21 cells, and in DMEM medium with 2% FBS. Briefly, 0.25× 10^6^/mL BHK21 cells were seeded in each well of a 12-well plate for 12h at 37°C to allow BHK21 cell adherence. Then, a serial dilution of each virus stock from YFV subculture in DMEM medium (without FBS, 1% penicillin/streptomycin, and 1% glutamine) was performed from 10^−1^ to 10^−7^. Then, 400 μL of each dilution was added in each well for virus adsorption 1-2hr. After this, the virus was discarded and each well was overlaid with 4mL 2% DMEM media containing final 0.8% complete LMP agarose. After 5 days of incubation at 37°C, the plaque visualization was made using 4% PFA fixation and 0.2% crystal violet in 20% ethanol staining to visualize the plaques.

### Human liver and brain tissue collection

Human liver tissue used in this study was collected from a patient who underwent surgical resection for hepatic hemangioma. Human brain tissue used in this study was collected from the unaffected area of a brain of a patient who died of epilepsy. The collection protocol was approved by the Institutional Review Board of Beijing Chao-Yang Hospital, Capital Medical University, and written informed consent regarding tissue donation was obtained from the patient and family before surgery.

### Human liver cell suspension preparation

The tissue sample was minced into small pieces and incubated in digestion buffer containing 1× PBS, collagenase D (1 mg ml−1, Sigma-Aldrich), dispase (2.4 mg ml−1, Thermo Fisher Scientific), DNase (0.2 mg ml−1, Sigma-Aldrich) and 3% heat-inactivated fetal bovine serum (FBS, Invitrogen) for 30 min at 37 °C. Cells suspensions were dissociated and passed through a 100 μm cell strainer (BD).

### Cell preparation for scRNA-seq

Thaw the cryovial containing frozen human liver single cells at 37℃, then transfer suspension to a centrifuge tube with RPMI1640 (Gibco 11875119)+10% FBS(Gibco 10100147C), centrifuged for 5 min at 300×g at 4°C. The cells were resuspended in RPMI1640+2% FBS. Cell count and viability was estimated using SeekMate Tinitan Fluorescence Cell Counter (SeekGene M002C) with AO/PI reagent after removal erythrocytes (Solarbio R1010) and then debris and dead cells removal was decided to be performed or not (Miltenyi 130-109-398/130-090-101). Finally fresh cells were washed twice in the RPMI1640 and then resuspended at 1×106 cells per ml in RPMI1640 and 2% FBS.

### Single cell RNA-seq library construction and sequencing

Single-cell RNA-Seq libraries were prepared using SeekOne® DD Single Cell 3’ library preparation kit (SeekGene Catalog No.K00202). Briefly, appropriate number of cells were mixed with reverse transcription reagent and then added to the sample well in SeekOne® chip S3. Subsequently, Barcoded Hydrogel Beads (BHBs) and partitioning oil were dispensed into corresponding wells separately in chip S3. After emulsion droplet generation reverse transcription were performed at 42℃ for 90 minutes and inactivated at 85℃ for 5 minutes. Next, cDNA was purified from broken droplet and amplified in PCR reaction. The amplified cDNA product was then cleaned, fragmented, end repaired, A-tailed and ligated to sequencing adaptor. Finally, the indexed PCR were performed to amplified the DNA representing 3’ polyA part of expressing genes which also contained Cell Barcode and Unique Molecular Index. The indexed sequencing libraries were cleanup with VAHTS DNA Clean Beads (Vazyme N411-01), analyzed by Qubit (Thermo Fisher Scientific Q33226) and Bio-Fragment Analyzer (Bioptic Qsep400). The libraries were then sequenced on GeneMind SURFSeq 5000 with PE150 read length.

### Single cell RNA-seq data analysis

scRNA-seq data processing and analysis were conducted using Seurat (v5.3.0). Quality control was applied by retaining cells that met the following criteria: nFeature_RNA between 200 and 5000, nCount_RNA below 20000, and mitochondrial gene content less than 10%. This filtering step resulted in a total of 8271 cells. The data were subsequently normalized and scaled using the standard Seurat workflow. Expression values of the genes of interest were then extracted to generate visualizations and compute mean expression levels.

### Data availability

ScRNA-seq raw data have been deposited in the Genome Sequence Archive (Genomics, Proteomics & Bioinformatics 2021) in National Genomics Data Center, China National Center for Bioinformation / Beijing Institute of Genomics, Chinese Academy of Sciences (GSA-Human: HRA013453).

### Statistical analysis

Unless otherwise specified, experiments were repeated at least three times. Bar graphs show mean ± SEM. Unpaired two-tailed t-tests were used for p-values unless otherwise noted.

## Extended Figures

**Extended Figure 1.**
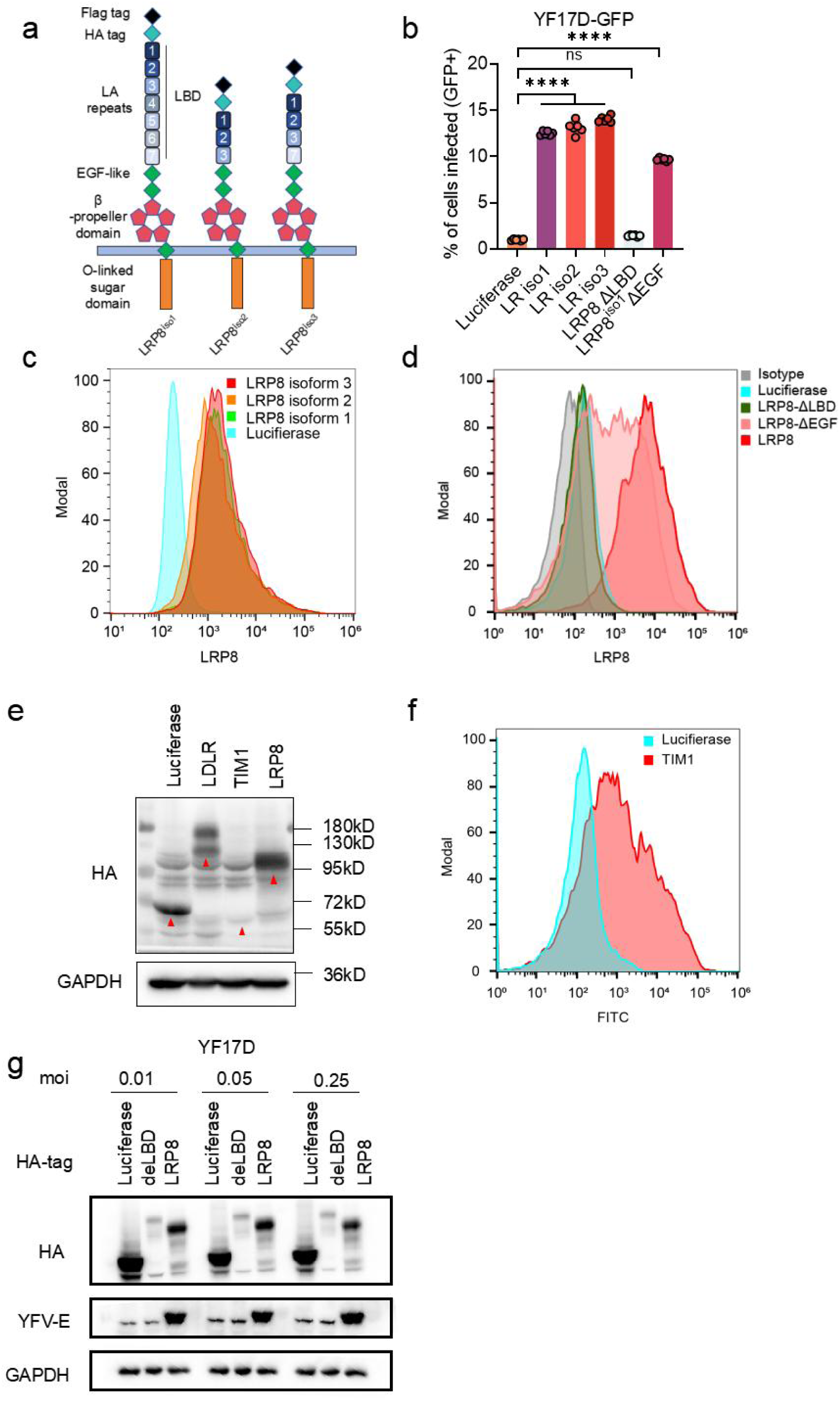
Different major isoforms of LRP8 have similar activities in promoting YF infection. **a,** Schematic structure of LRP8 isoforms; **b,** In HEK293T cells, luciferase, LRP8 isoforms and its truncations were transiently overexpressed and the cells are infected by YFV 17D pseudoviruses. The ratios of infected cells were analyzed using FACS at 24hpi. **c,d,** Expression of the isoforms and its truncations on cell surface were determined with anti-LRP8 by FACS. **e, f**, Luciferase, LDLR, LRP8 or TIM1 containing 3xHA tag were analyzed by western blot in HEK293T cells or FACS. **g**. In HEK293T cells, luciferase, LRP8-ΔLBD and LRP8 were transiently overexpressed and the cells are infected by live YF17D with different MOI for 48h. The infected cells were analyzed using Western blots. Statistical analysis was performed using one-way analysis of variance (ANOVA) with Dunnett’s multiple comparisons test(b). The statistics shown are Mean ± SEM. *: P < 0.05; **: P < 0.01; ***: P < 0.001; ****: P < 0.0001, ns, not significant. The replicate in the figure indicates biological replicates, the experiments were repeated at least twice.

**Extended Figure 2.**
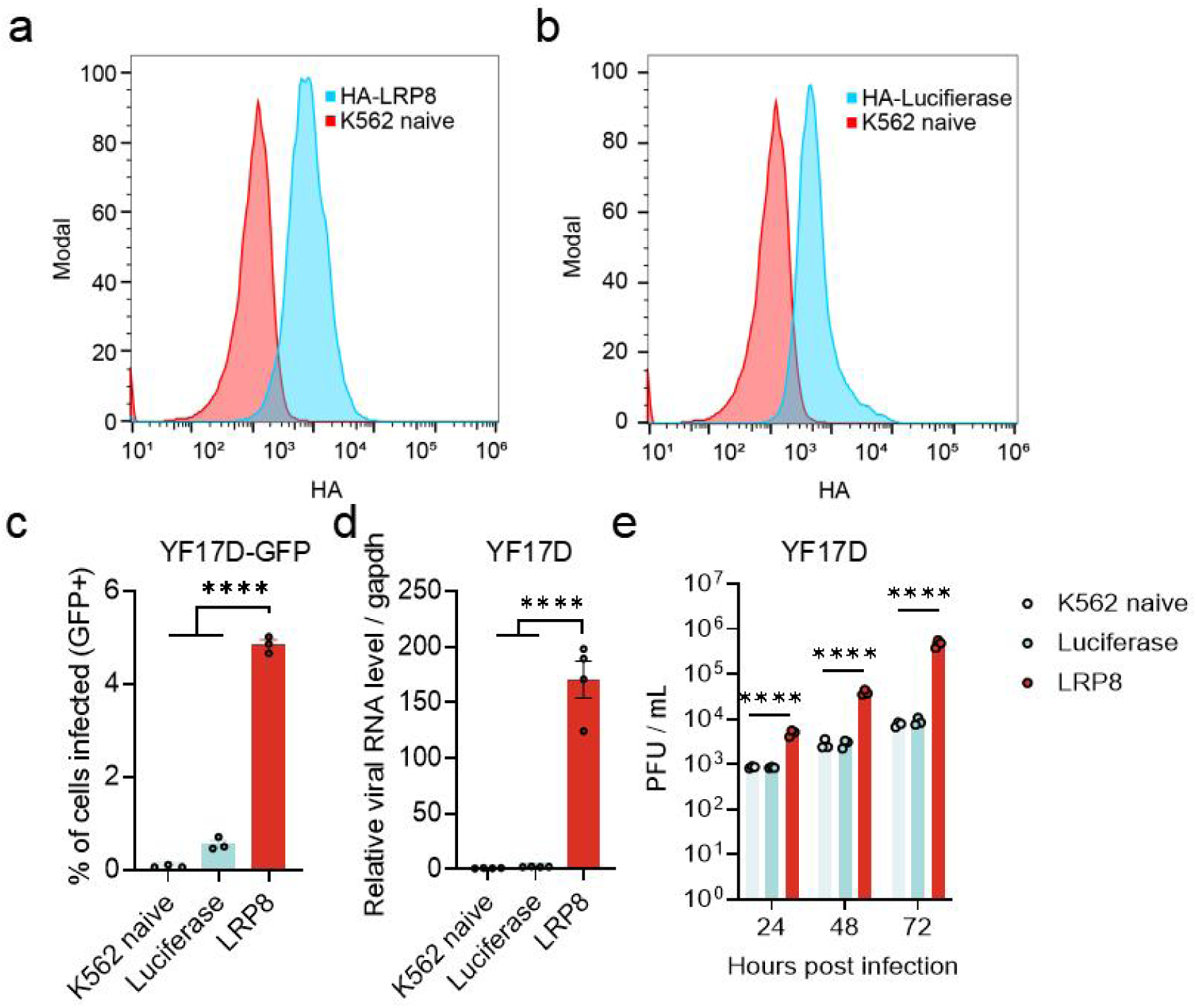
LRP8 promotes YF17D infection in K562 cells. **a, b**. LRP8 and luciferase expression on K562 cell surface were determined by FACS with HA antibody staining. **c**. In K562 cells, HA-tagged luciferase and LRP8 were stably overexpressed and the cells are infected by YFV 17D pseudoviruses and the infection rates were determined by FACS at 24 hpi. **d**. In K562 cells, HA-tagged luciferase and LRP8 were stably overexpressed and the cells are infected by live YFV 17D and viral RNA level in cells were determined by qRT-PCR at 48 hpi. **e**. Infectious viral particles from d were determined by plaque assay. Statistical analysis was performed using two-way analysis of variance (ANOVA) with Dunnett’s multiple comparisons test(c-e). The statistics shown are Mean ± SEM. *: P < 0.05; **: P < 0.01; ***: P < 0.001; ****: P < 0.0001, ns, not significant. The replicate in the figure indicates biological replicates, the experiments were repeated at least twice.

**Extended Figure 3.**
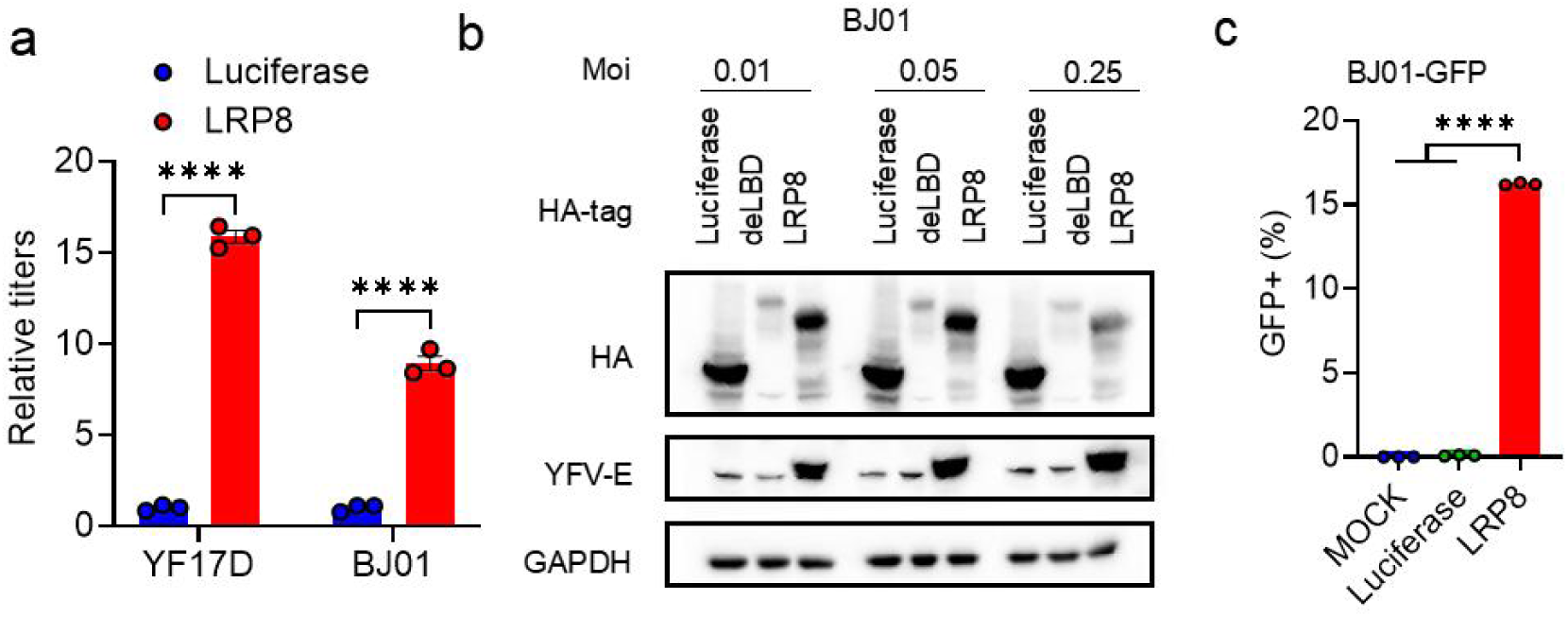
LRP8 promotes the infection by live viruses of YF17D, BJ01 and BJ01-GFP pseudovirus. **a**. Supernatant of HEK293T expressing Luciferase and LRP8 infected with YF17D (0.05 MOI) and BJ01(0.05 MOI). At 48hpi, the viral titers were determined by plaque assay in BHK21 cells. **b**. In HEK293T cells, luciferase, LRP8-ΔLBD and LRP8 were transiently overexpressed and the cells are infected by live BJ01 with different MOI for 48h. The infected cells were analyzed using Western blots. **c**. In HEK293T cells, luciferase and LRP8 were transiently overexpressed and the cells are infected by YFV BJ01 pseudoviruses. The ratios of infected cells were analyzed using FACS at 24 hpi. Statistical analysis was performed using two-way analysis of variance (ANOVA) with Uncorrected Fisher’s LSD test (a), Dunnett’s multiple comparisons test(c). The statistics shown are Mean ± SEM. *: P < 0.05; **: P < 0.01; ***: P < 0.001; ****: P < 0.0001, ns, not significant. The replicate in the figure indicates biological replicates, the experiments were repeated at least twice.

**Extended Figure 4.**
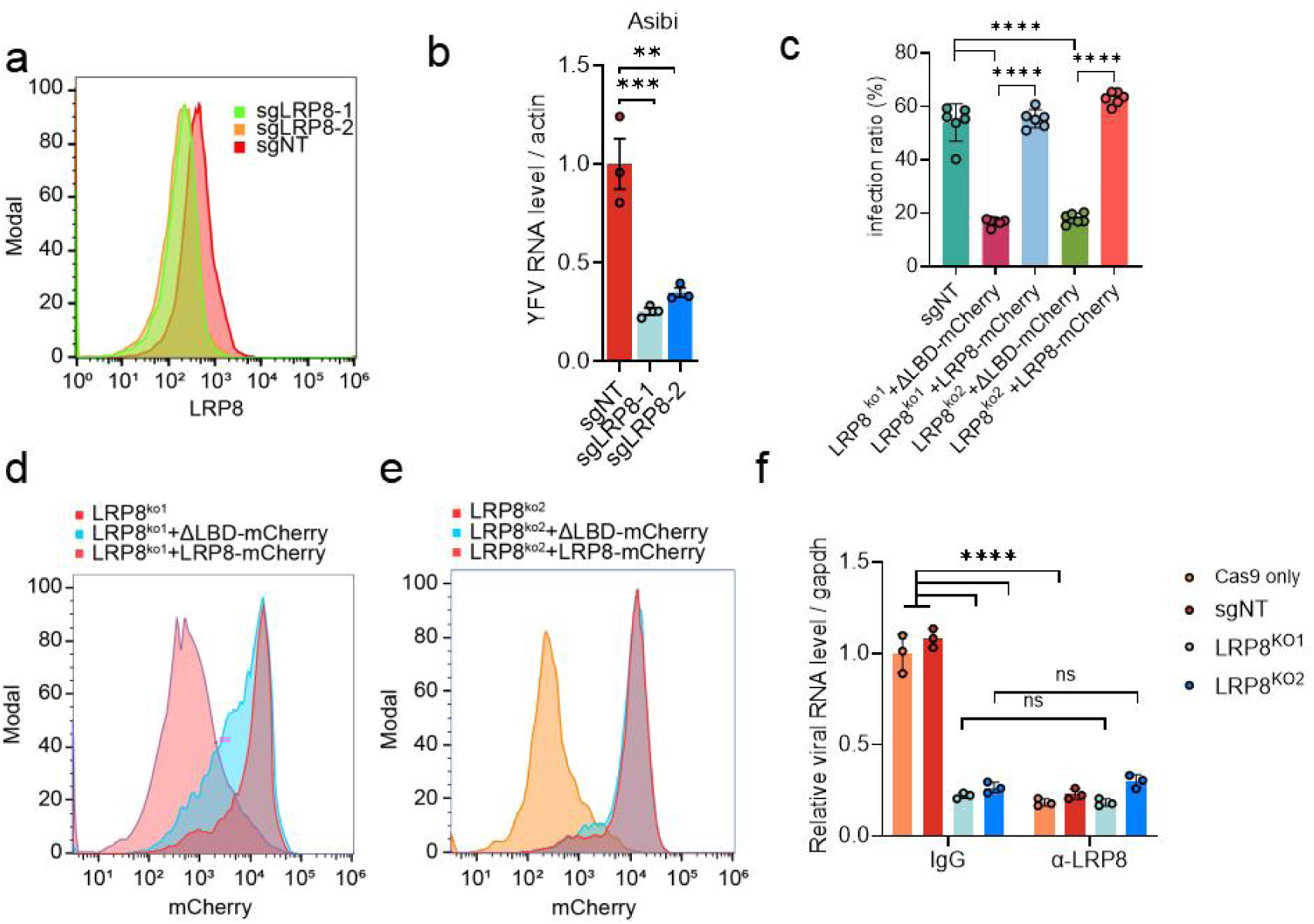
Deficiency of LRP8 decreases live Asibi and YF17D infection. **a.** U87MG cells expressing Cas9/sgLRP8 single clones and Cas9 only were tested by FACS with anti-LRP8. **b**, Knockout cells were infected with the clinical Asibi (0.2 moi), viral RNA was quantified by RT–qPCR. **c-e**, In two LRP8 KO cells, ΔLBD-mCherry and LRP8-mcherry plasmids were electroporated and 48h later the cells are infected with YF17D. At 48 hpi, the cells were stained for YFV E protein and quantitated by fluorescence microscope. ΔLBD-mCherry and LRP8-mcherry expressions were determined by FACS. **f**. U87MG-sgLRP8 and control U87MG cells expressing only Cas9 were preincubated with 10 μg/ml anti-LRP8 (sc-1A1) or IgG before being infected by YF17D. At 48 hpi, the infections were analyzed by qRT-PCR. Statistical analysis was performed using two-way analysis of variance (ANOVA) with Turkey’s multiple comparisons test (b, c, f). The statistics shown are Mean ± SEM. *: P < 0.05; **: P < 0.01; ***: P < 0.001; ****: P < 0.0001, ns, not significant. The replicate in the figure indicates biological replicates, the experiments were repeated at least twice.

**Extended Figure 5.**
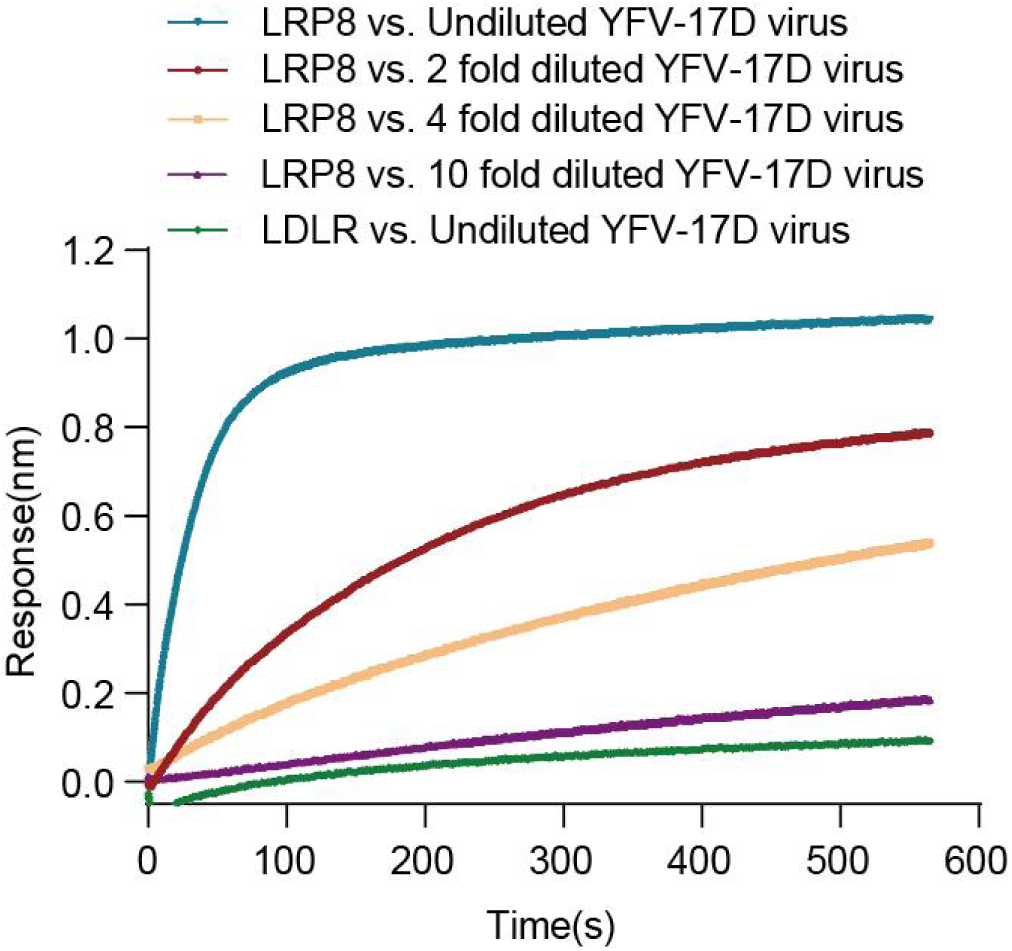
Purified LRP8 LBD or LDLR LBD were biotinated and immobilized on the probe of biolayer inferometry (BLI). The binding of YFV 17D viral particles to LRP8 or LDLR was measured by BLI assay.

**Extended Figure 6.**
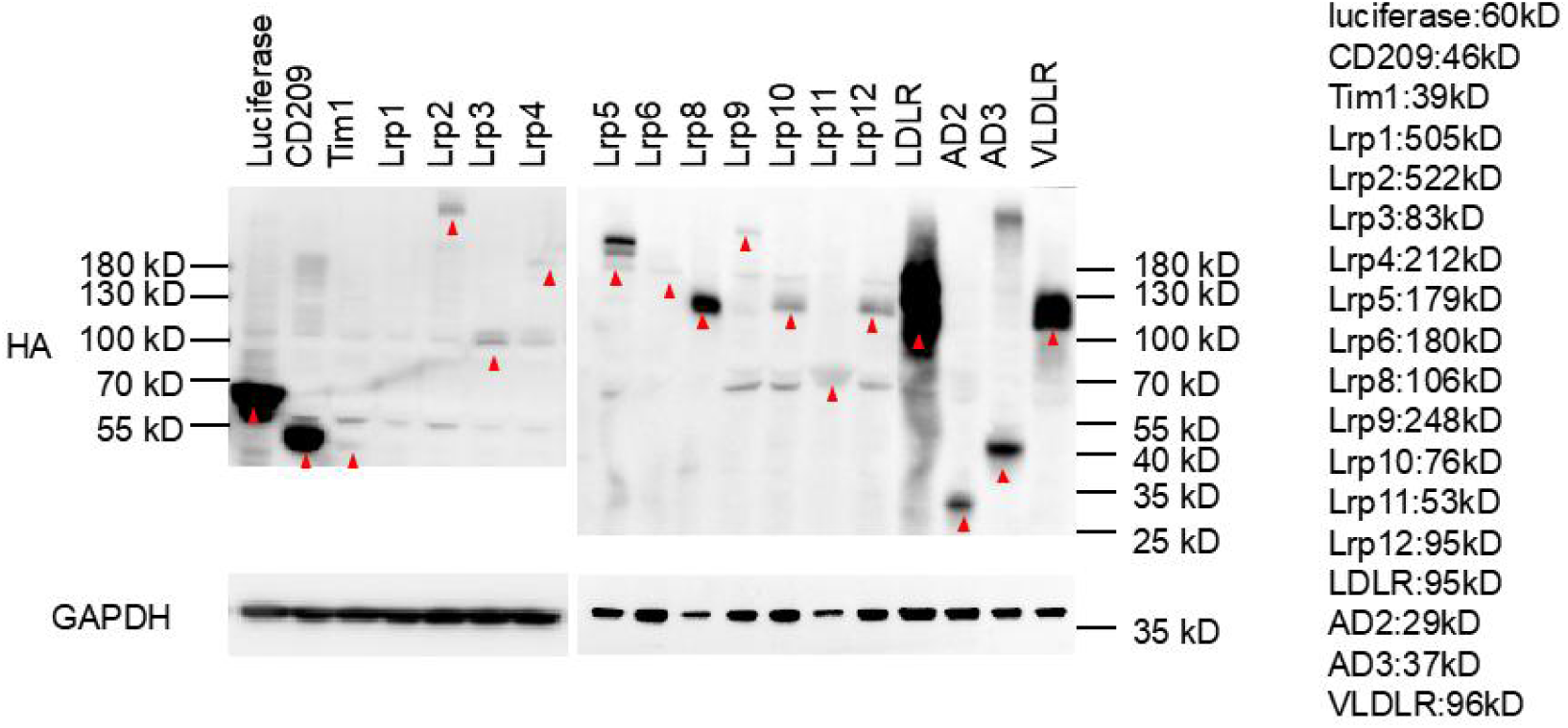
Expression of different members of the LDLR family. In HEK293T cells, plasmids expressing different LDLR family members were transfected into HEK293T for 48 hours and the target protein containing 3X HA tag were analyzed by Western blot. The corresponding molecular weight of each protein were marked at the right of figure.

**Extended Figure 7.**
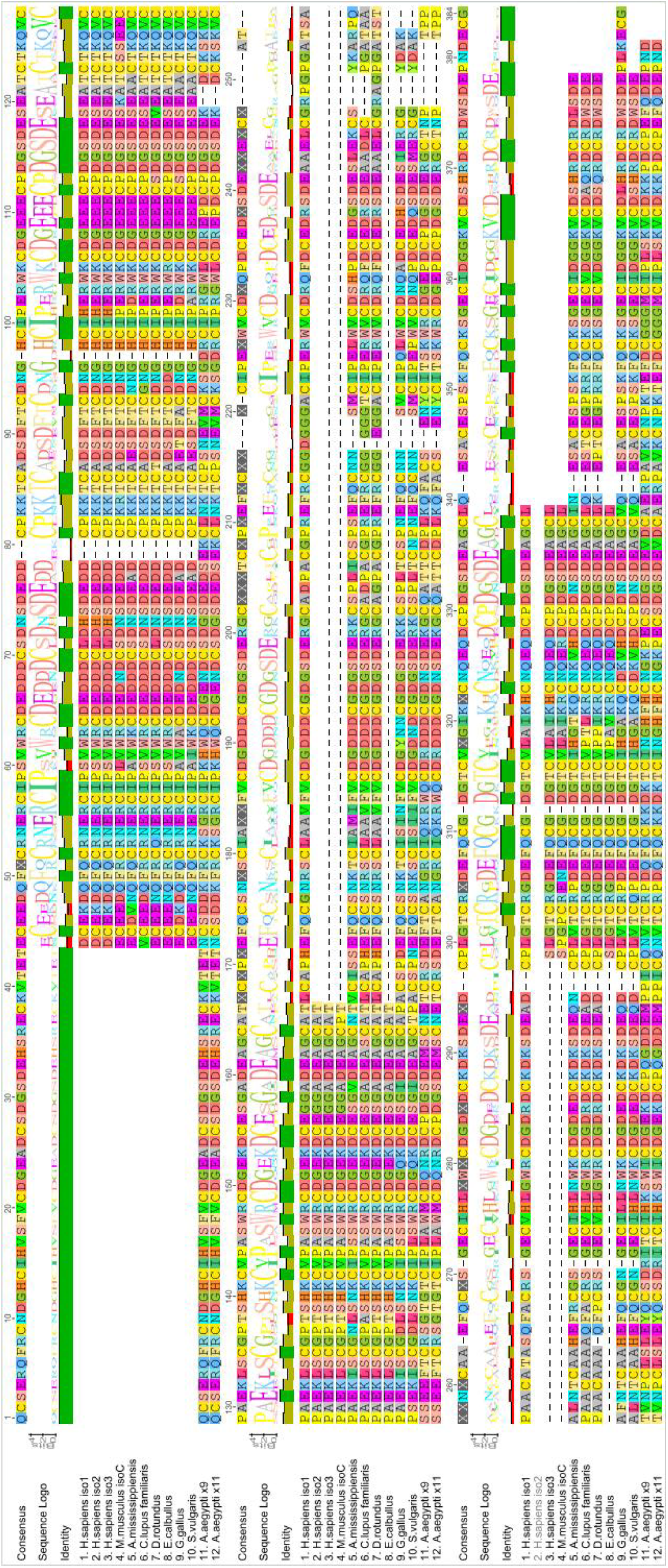
The amino acid sequence alignment of LRP8 ligand binding domain (LBD). The LRP8 LBD amino acid sequences of human, mouse, mosquito, A. mississippiensis, bat, chicken, bird, dog, horse, green monkey were downloaded from NCBI and aligned using Geneious software.

**Extended Figure 8.**
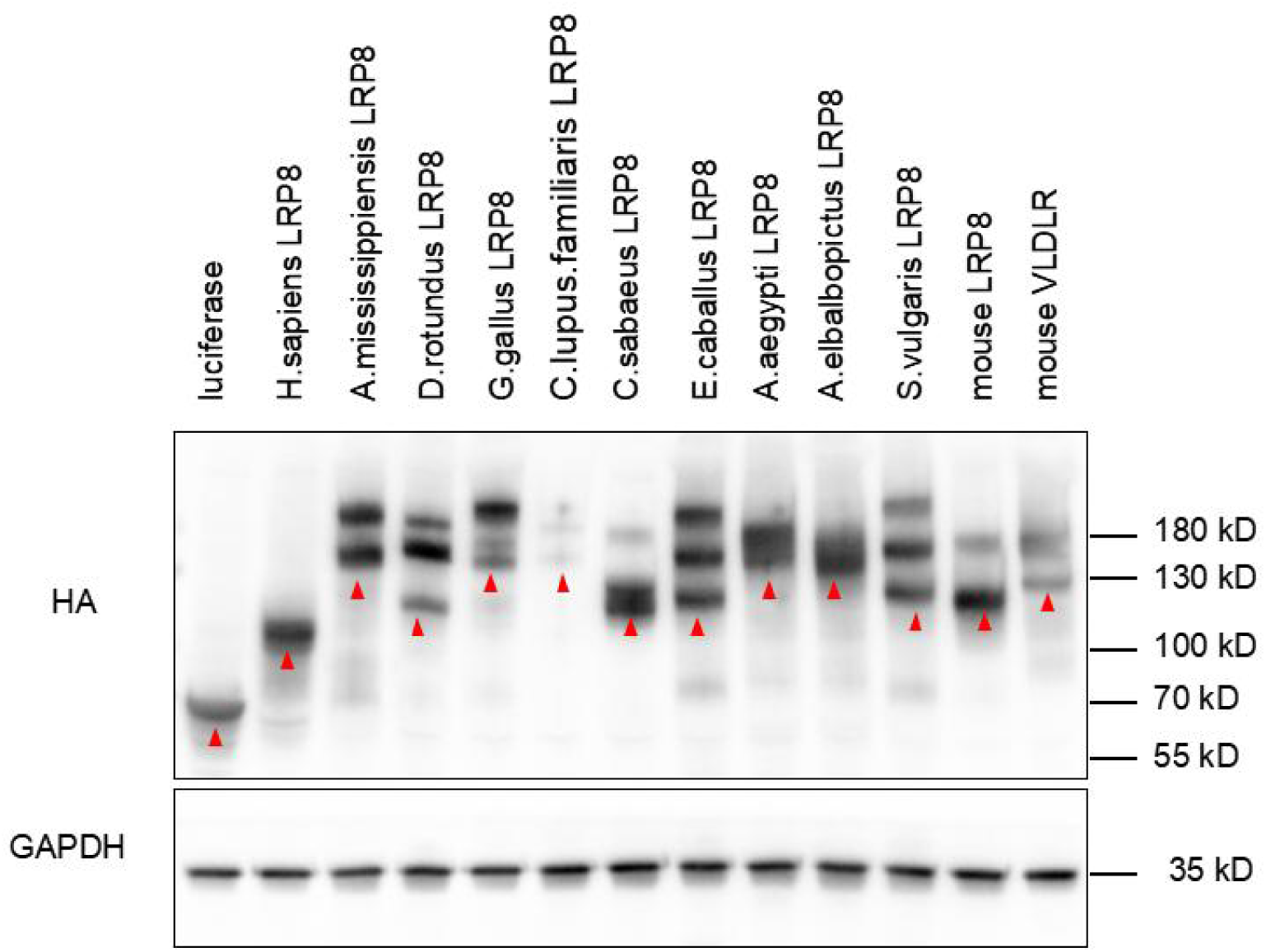
Expression of LRP8 homologs from different species. In HEK293T cells, plasmids expressing different LRP8 orthologs were transfected into HEK293T for 48 hours and the target protein containing 3X HA tag were analyzed by Western blot.

**Extended Figure 9.**
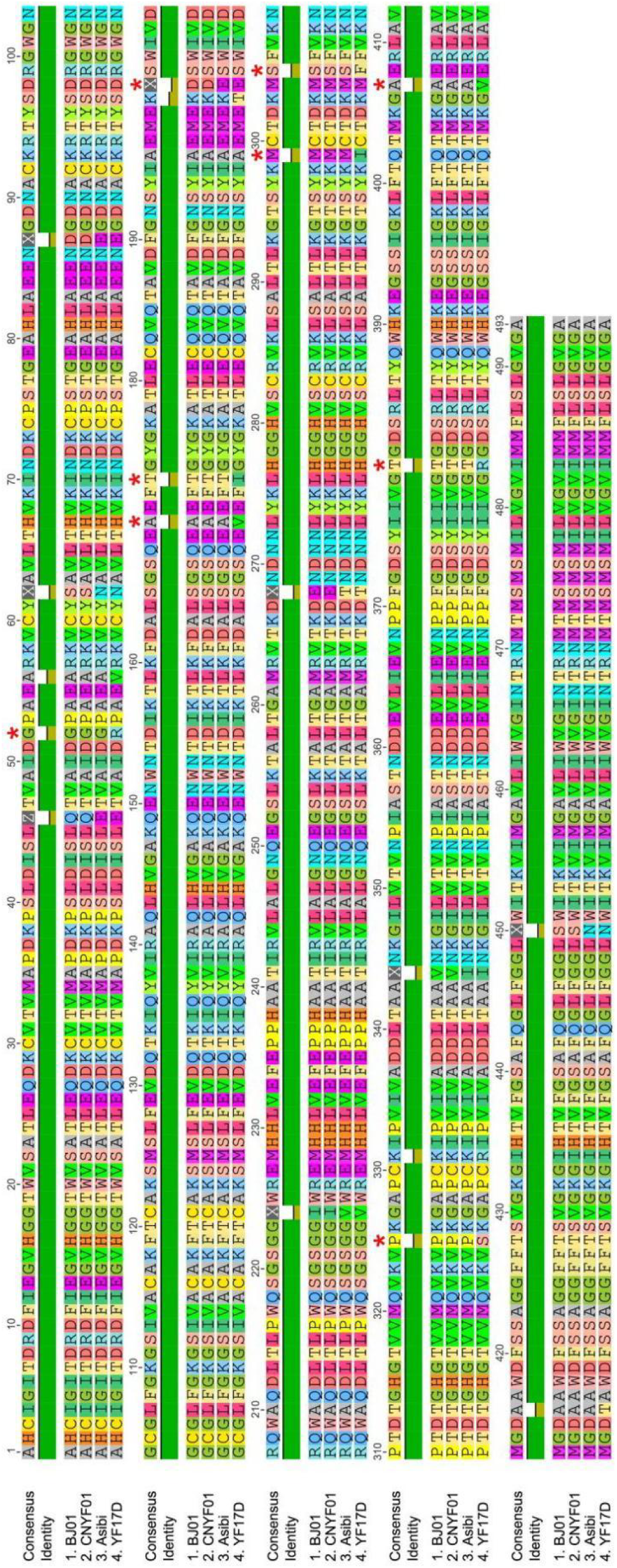
The amino acid sequence alignment of YF strain envelope proteins. The amino acid sequence of the envelope protein of YF BJ01, CNYF01, Asibi, YF17D strains were downloaded from NCBI and aligned using Geneious software. Asterisks indicate sites of mutations in E.

**Extended Figure 10.**
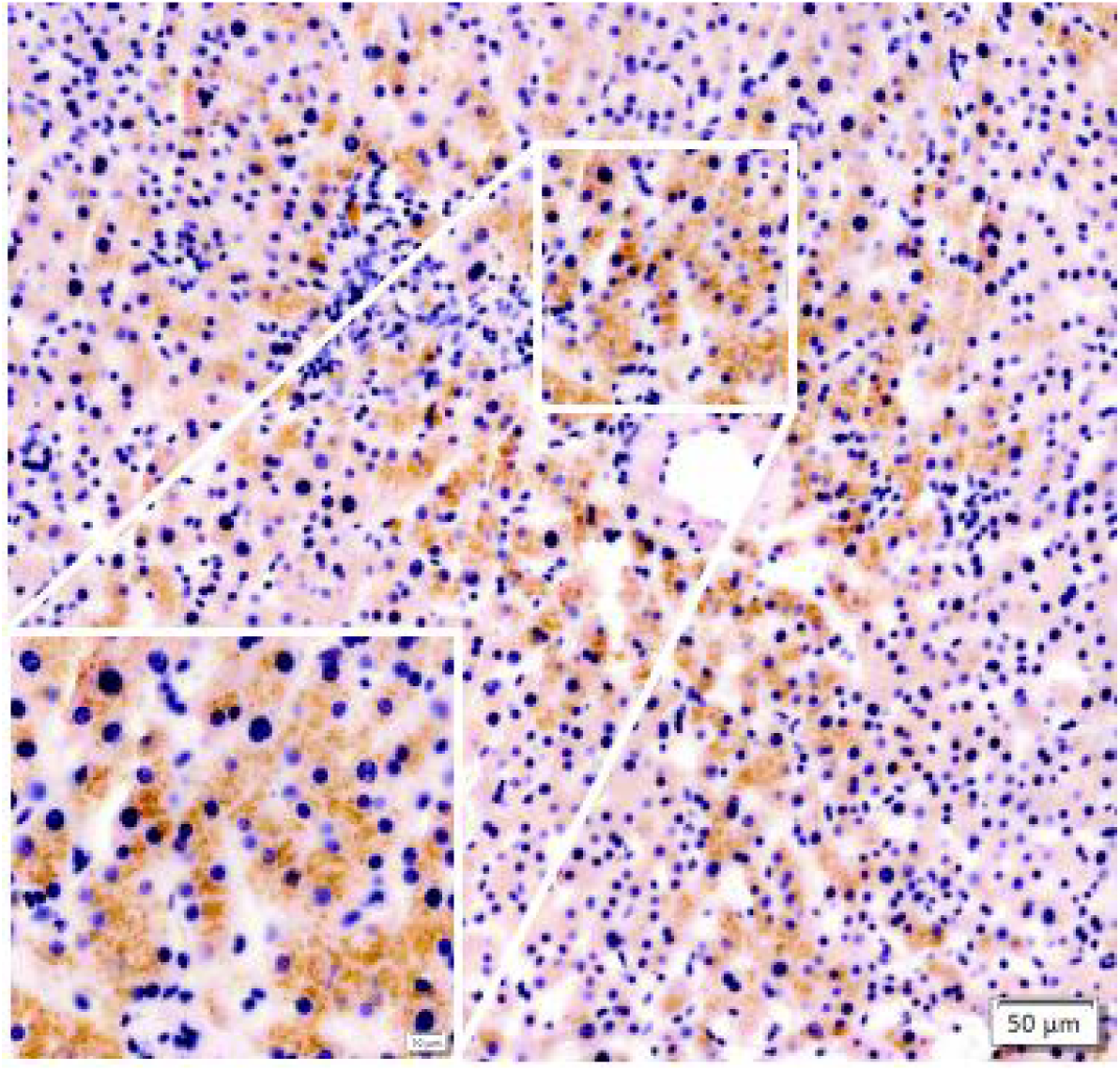
LRP8 expression were determined in human liver sample by immunohistochemistry staining with a LRP8 antibody (Thermo 3H2 clone).

**Extended Figure 11.**
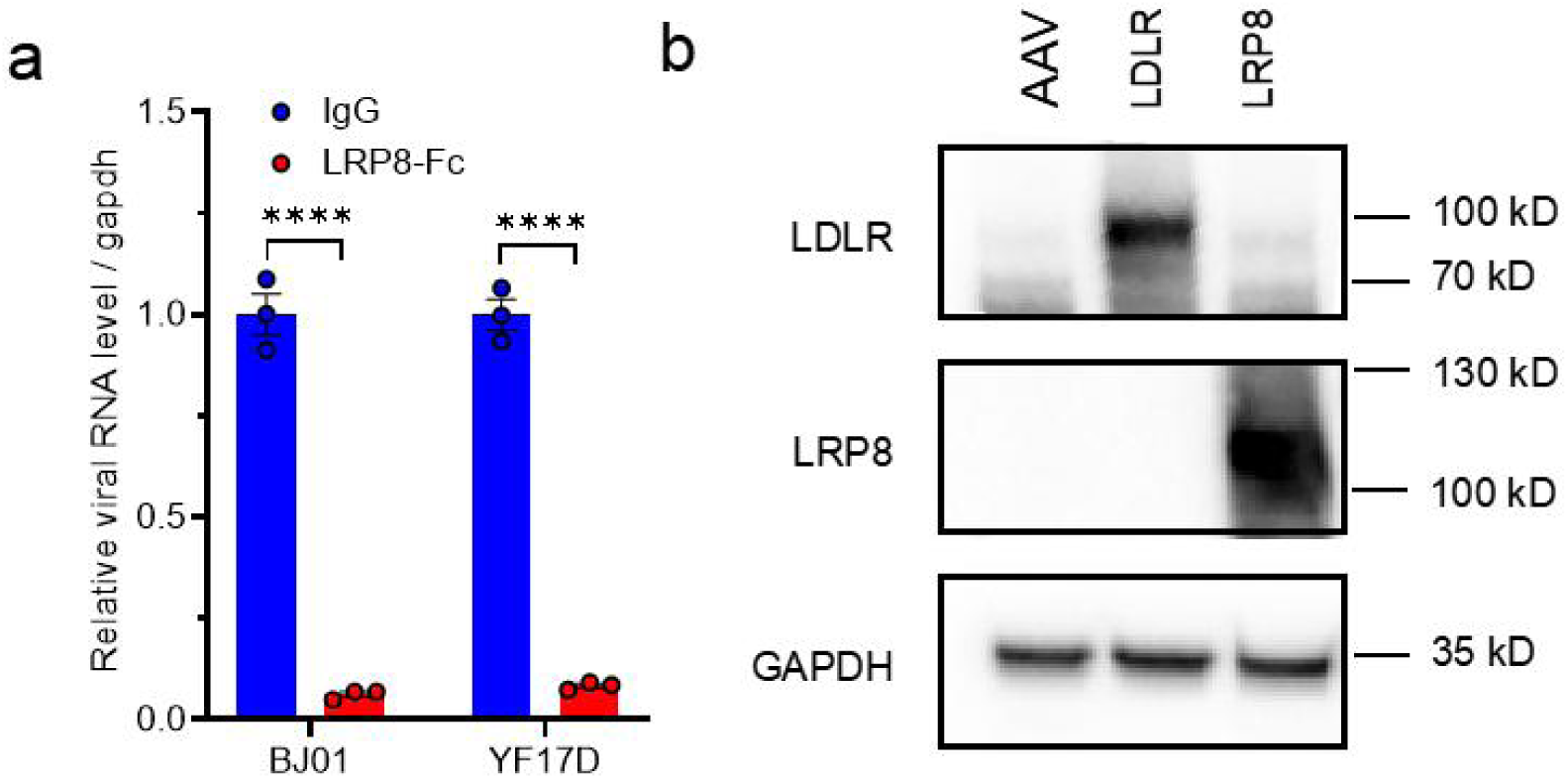
Soluble LRP8-Fc blocks BJ01 and YF17D infection and validation of the expression of LDLR and LRP8 delivered by AAV in mouse liver. **a**. Preincubation of soluble LRP8 (200 μg/ml) with 0.5 MOI of BJ01 and Asibi for 1 hour, and the protein-virus mixture were added into PHHs for 48 hours. The viral RNA was quantified by qRT-PCR (three biological repeats). **b**. hLDLR and hLRP8 in mouse liver were determined at 3 weeks post AAV delivering by Western blot. Statistical analysis was performed using two-way analysis of variance (ANOVA) with Uncorrected Fisher’s LSD test (a). The statistics shown are Mean ± SEM. *: P < 0.05; **: P < 0.01; ***: P < 0.001; ****: P < 0.0001, ns, not significant. The replicate in the figure indicates biological replicates, the experiments were repeated at least twice.

**Extended Figure 12.**
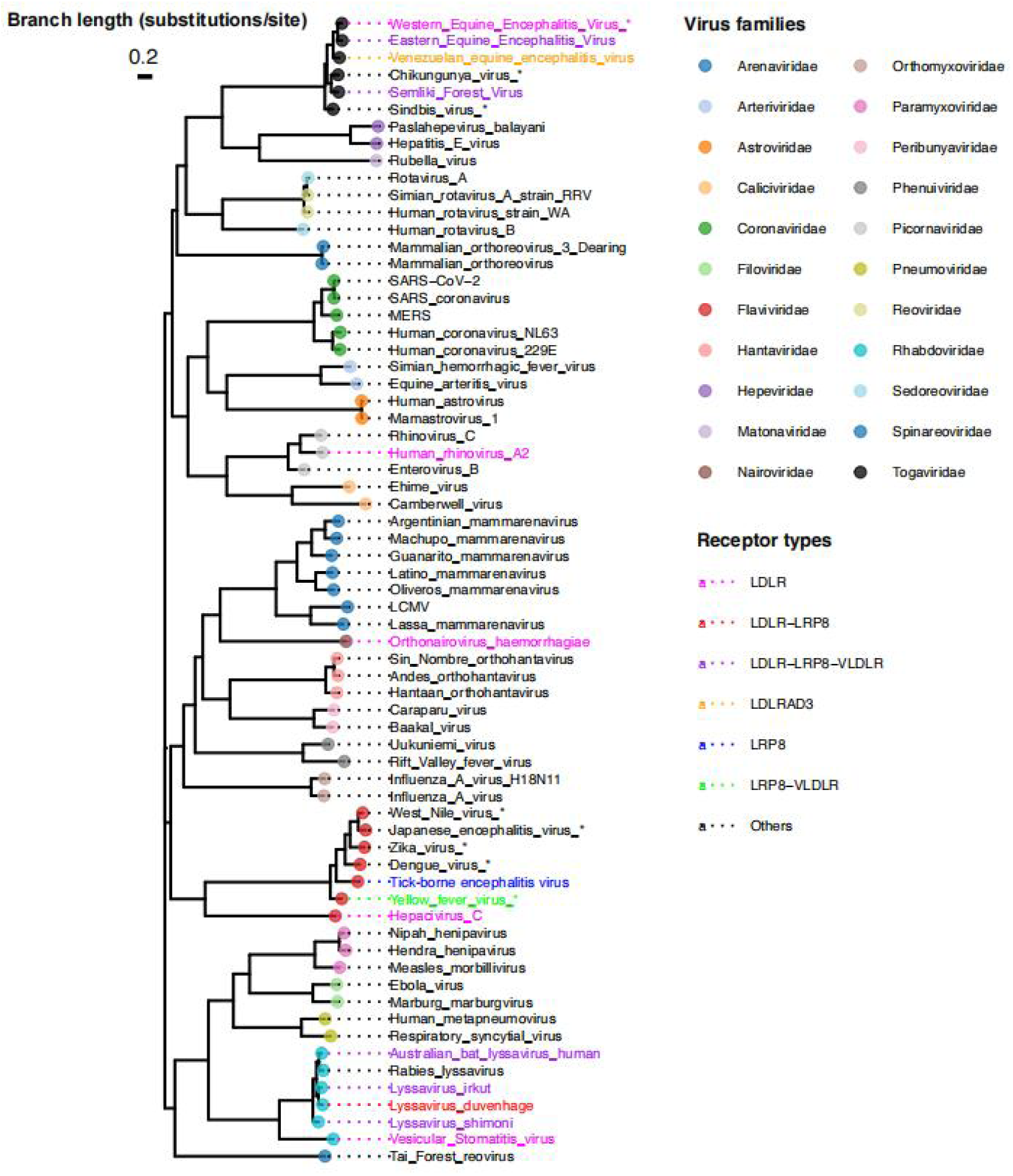
Phylogenetic Tree Construction of RNA Virus RdRPs. To investigate the evolutionary relationships of RNA-dependent RNA polymerases (RdRPs) among human-infecting RNA viruses, we curated 24 RNA virus families from the Virus-Host Database. Receptor-annotated viruses were identified via the ViralReceptor database, covering 17 families. RdRP protein sequences were collected from NCBI, GenBank, and UniProtKB/Swiss-Prot. Among 94 viruses, all except Hepatitis delta virus, Echovirus E8, and Retroviridae had identifiable RdRPs, yielding 22 families for analysis. Sequences were aligned using MAFFT (v7.526) with the --localpair --maxiterate 1000 options, and a maximum likelihood tree was built with FastTree (v2.1.11). Visualization was done using ggtree (v3.21) in R (v4.4.3), highlighting families with known human receptors. Asterisks indicate mosquito-borne viruses.

## Notes

### Competing Interest Statement

The authors have declared no competing interest.

### Summary of Updates

Authorship changes, two authors were removed from the author list and were acknowledged instead.

